# Cryptic inoviruses are pervasive in bacteria and archaea across Earth’s biomes

**DOI:** 10.1101/548222

**Authors:** Simon Roux, Mart Krupovic, Rebecca A. Daly, Adair L. Borges, Stephen Nayfach, Frederik Schulz, Jan-Fang Cheng, Natalia N. Ivanova, Joseph Bondy-Denomy, Kelly C. Wrighton, Tanja Woyke, Axel Visel, Nikos C. Kyrpides, Emiley A. Eloe-Fadrosh

**Affiliations:** DOE Joint Genome Institute, Walnut Creek, CA 94598, USA; Institut Pasteur, Unité Biologie Moléculaire du Gène chez les Extrêmophiles, Paris, 75015, France; Department of Soil and Crop Sciences, Colorado State University, Fort Collins, CO 80521, USA; Department of Microbiology and Immunology, University of California, San Francisco, San Francisco, CA 94143, USA; Quantitative Biosciences Institute, University of California, San Francisco, San Francisco, CA 94143, USA

**Keywords:** virus, bacteriophage, archaeal virus, viral diversity, chronic virus, host-pathogen interaction

## Abstract

Bacteriophages from the *Inoviridae* family (inoviruses) are characterized by their unique morphology, genome content, and infection cycle. To date, a relatively small number of inovirus isolates have been extensively studied, either for biotechnological applications such as phage display, or because of their impact on the toxicity of known bacterial pathogens including *Vibrio cholerae* and *Neisseria meningitidis*. Here we show that the current 56 members of the *Inoviridae* family represent a minute fraction of a highly diverse group of inoviruses. Using a new machine learning approach leveraging a combination of marker gene and genome features, we identified 10,295 inovirus-like genomes from microbial genomes and metagenomes. Collectively, these represent six distinct proposed inovirus families infecting nearly all bacterial phyla across virtually every ecosystem. Putative inoviruses were also detected in several archaeal genomes, suggesting that these viruses may have occasionally transferred from bacterial to archaeal hosts. Finally, we identified an expansive diversity of inovirus-encoded toxin-antitoxin and gene expression modulation systems, alongside evidence of both synergistic (CRISPR evasion) and antagonistic (superinfection exclusion) interactions with co-infecting viruses which we experimentally validated in a *Pseudomonas* model. Capturing this previously obscured component of the global virosphere sparks new avenues for microbial manipulation approaches and innovative biotechnological applications.

## Introduction

Inoviruses, bacteriophages from the *Inoviridae* family, exhibit unique morphological and genetic features. While the vast majority of known bacteriophages carry double-stranded DNA (dsDNA) genomes encapsidated into icosahedral capsids, inoviruses are instead characterized by rod-shaped or filamentous virions which carry a circular single-stranded DNA (ssDNA) genomes of ~5-20kb^1–3^. One of the most striking features of inoviruses is their ability to establish a chronic infection whereby the viral genome resides within the cell in either exclusively episomal state or integrated into the host chromosome and virions are continuously released without killing the host^1,2,4^ (Figure 1A). Owing to their unique morphology and simple genome amenable to genetic engineering, several inoviruses are widely used for biotechnological applications, including phage display or as drug delivery nanocarriers^5–8^.

**Figure 1.**
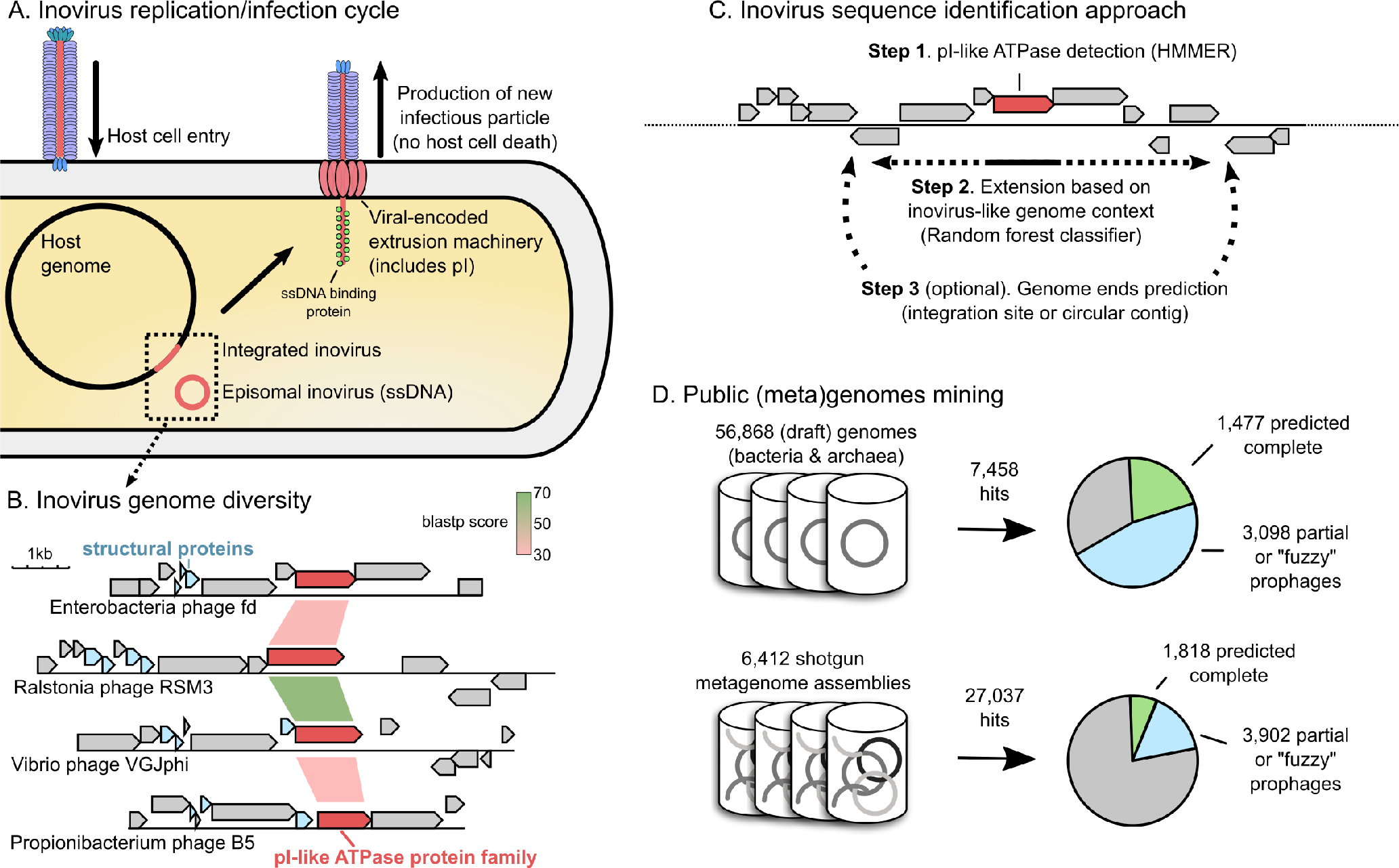
Overview of inovirus infection cycle, diversity, and sequence detection process. A. Schematic of inovirus persistent infection cycle and virion production. Inovirus genomes and particles are not to scale relative to the host cell and genome. B. Comparison of selected inovirus genomes from isolates. The pI-like genes (the most conserved genes) are colored in red, and sequence similarity between these genes (based on blastp) is indicated with colored links between genomes. Putative structural proteins that can be identified based on characteristic features (gene length and presence of a transmembrane domain) are colored in blue. C. Representation of the new inovirus detection approach. D. Results of the search for inovirus sequences in prokaryote genomes and assembled metagenomes, after exclusion of putative false-positive through manual inspection of predicted pI proteins (see Supplementary Text). Predictions for which genome ends could be identified are indicated in green, while predictions without clear ends, i.e. partial genomes or “fuzzy” prophages with no predicted attachment site, are in blue, adding up to 10,295 curated predictions in total. Sequences for which no inovirus genome could be predicted around the initial pI-like gene are in gray. See also Figure S1 & S2.

Ecologically, the chronic infection cycle of inoviruses results in extended viral residence time in the host cell, enabling inoviruses to continuously manipulate and alter their host’s phenotype. Although cultivated inoviruses are known to infect hosts from only 5 bacterial phyla and 10 genera, some are already known to confer or modulate bacterial pathogenicity^9^, while others can increase the growth rate of their host (conditional mutualism)^10^. For instance, an inovirus prophage, CTXphi, encodes and expresses the major virulence factor of toxigenic *Vibrio cholerae*^11,12^. In other bacterial hosts including *Pseudomonas*, *Neisseria*, and *Ralstonia*, inovirus infections indirectly influence pathogenicity by altering biofilm formation and host colonization abilities^9,13–16^.

Despite these remarkable properties, their elusive life cycle and peculiar genomic and morphological properties have hampered systematic discovery of additional inoviruses: to date, only 56 inovirus genomes have been described^4^. Most inoviruses do not elicit negative effects on the growth of their hosts when cultivated in the laboratory and can thus easily evade detection. Furthermore, established computational approaches for detection of virus sequences in whole genome shotgun sequencing data are not efficient for inoviruses because of their unique and diverse gene content^17–19^ (Figure 1B). Finally, inoviruses are likely undersampled in viral metagenomes due to their long, flexible virions with low buoyant density^20,21^.

Here we unveil a substantial diversity of 10,295 inovirus sequences, derived from a broad range of bacterial and archaeal hosts. These were identified through an exhaustive search of 56,868 microbial genomes and 6,412 shotgun metagenomes using a new computational approach to identify known and novel putative inovirus genomes. These viruses likely represent a reclassified viral order composed of at least 6 families and 212 subfamilies, and encode a rich and mostly novel gene content. A large fraction of this inovirus-encoded genetic diversity is seemingly dedicated to manipulation of and maintenance in host populations, as well as interactions with co-infecting viruses. Overall, these data clearly indicate that inoviruses are far more widespread, diverse, and ecologically pervasive than previously appreciated, providing a robust foundation to further characterize their biology across multiple hosts and environments.

## Results

### Inoviruses are highly diverse and globally prevalent

To evaluate the global diversity of inoviruses, an analysis of all publicly available inovirus genomes was first conducted to identify characteristic traits that would enable discovery of new inovirus sequences (Table S1). Across the 56 known *Inoviridae* genomes, the gene coding for the morphogenesis (pI) protein, an ATPase of the FtsK-HerA superfamily, represented the only conserved marker gene (Figure 1A & B, Figure S1A). However, three additional features specific of inovirus genomes could be defined: (i) short structural proteins (30 to 90 aa) with a single predicted transmembrane domain (TMD; Table S1), (ii) genes either functionally uncharacterized or similar to other inoviruses, and (iii) shorter genes compared to those in typical bacterial or archaeal genomes (Figure S1B). These features were used to train a random forest classifier which, associated with the detection of a pI-like protein, was able to identify inovirus sequences from background host genome with 92.5% recall and 99.8% precision on our manually curated reference set (Figure 1C, Figure S1C, Supplementary Text).

This new detection approach was applied to 56,868 bacterial and archaeal genomes and 6,412 metagenomes publicly available from the IMG database^22^ (Table S2). After manual curation of edge cases and removal of detections not based on a clear inovirus-like ATPase, a total of 10,295 sequences were recovered (Figure 1D, Figure S2, Supplementary Text). From these, 5,964 distinct species were identified using genome-wide Average Nucleotide Identity (ANI), and only 38 of these included isolate inovirus genomes. About one third of these species (30%) encoded an “atypical” morphogenesis gene, with an N-terminal instead of C-terminal TMD (Figure S2). Although this atypical domain organization has been observed in four isolate species currently classified as inoviruses, some of these inovirus-like sequences might eventually be considered as entirely new groups of viruses. Sequence accumulation curves did not reach saturation, highlighting the large diversity of inoviruses yet to be sampled (Figure S1D).

Inovirus sequences were identified in 6% of bacterial and archaeal genomes (3,609 of 56,868), and 35% of metagenomes (2,249 of 6,412). More than half of the species (n=3,675) were exclusively composed of sequences assembled from metagenomes. These revealed that inoviruses are found in every major microbial habitat whether aquatic-, soil-, or human-associated, and throughout the entire globe (Figure 2 & Supplementary Text). Hence, inoviruses are much more diverse than previously estimated and globally distributed.

**Figure 2.**
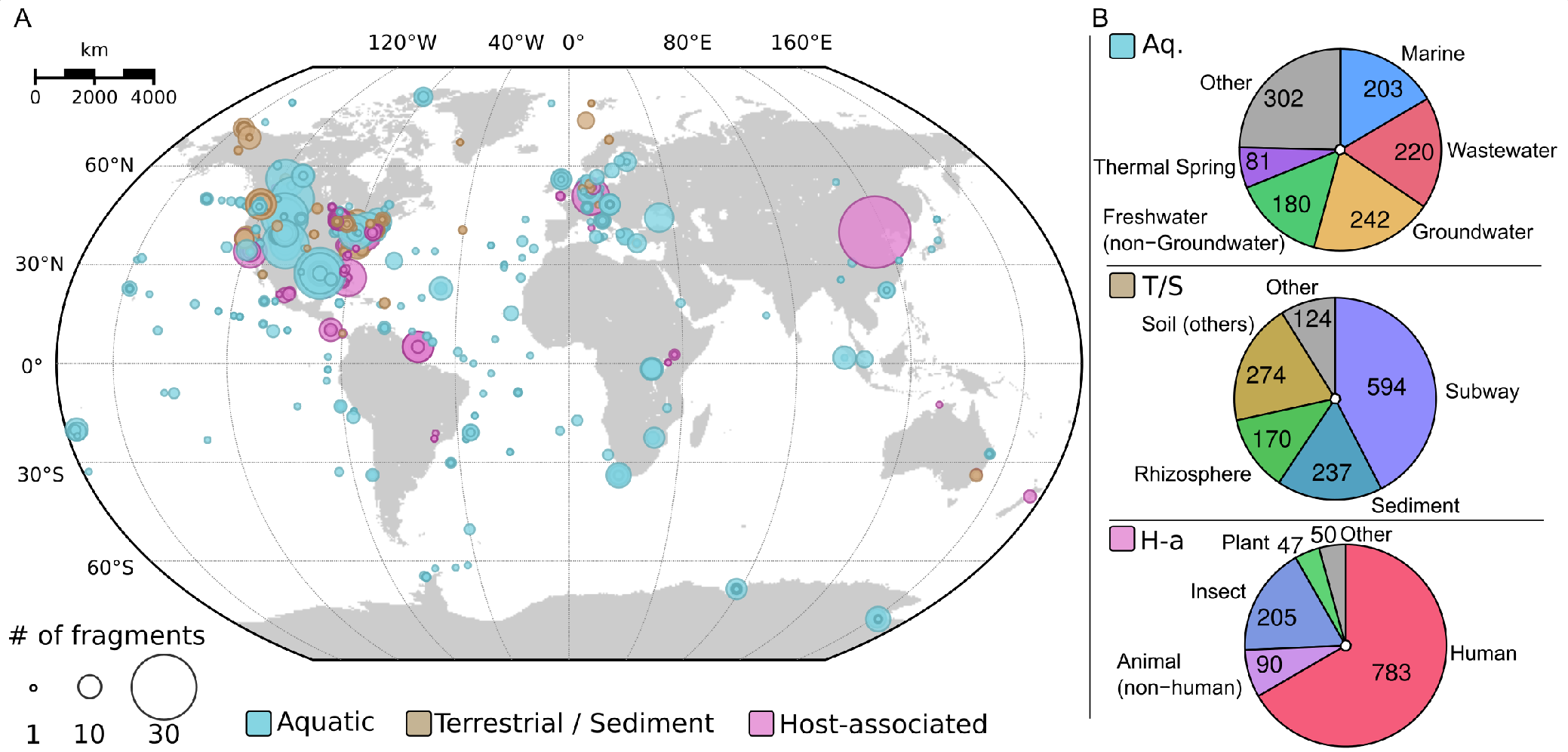
Geographic and biome distribution of inovirus sequences detected in metagenomes. A. Repartition of samples for which ≥ 1 inovirus sequence(s) were detected. Each sample is represented by a circle proportional to the number of inovirus detections, and colored according to their ecosystem type. B. Breakdown of the number of inovirus detections by ecosystem subtype for each major ecosystem. Aq.: Aquatic, T/S: Terrestrial / Sediment, H-a: Host-associated.

### Inoviruses infect a broad diversity of bacterial hosts

To examine the host range of these newly discovered inoviruses, we focused on the 2,284 inovirus species directly associated with a host, i.e. proviruses derived from a microbial genome (Figure 3). The majority (90%) of these species were associated with Gamma- and Beta-proteobacteria, from which most known inoviruses were previously isolated (Table S1). The range of host genera within these groups was, however, vastly expanded, including clinical and ecologically relevant microbes such as *Azotobacter*, *Haemophilus*, *Kingella*, or *Nitrosomonas* (Table S3). The remaining 412 species strikingly increased the potential host range of inoviruses to 22 additional phyla, including the Candidate Phyla Radiation (CPR, Figure 3). For 3 of these (Acidobacteria, Chlamydiae, and Spirochaetes), only short inovirus contigs were detected, lacking host flanking regions which would provide confident host linkages. Hence, these contigs could potentially derive from sample contamination (e.g. from reagents), and inovirus presence within these phyla remains uncertain (Table S4). The notable host expansion is consistent with reported experimental observations of filamentous virus particles induced from a broad range of bacteria, e.g. *Mesorhizobium*, *Clostridium*, *Flavobacterium*, *Bacillus*, and *Arthrobacter*^23,24^(Figure 3).

**Figure 3.**
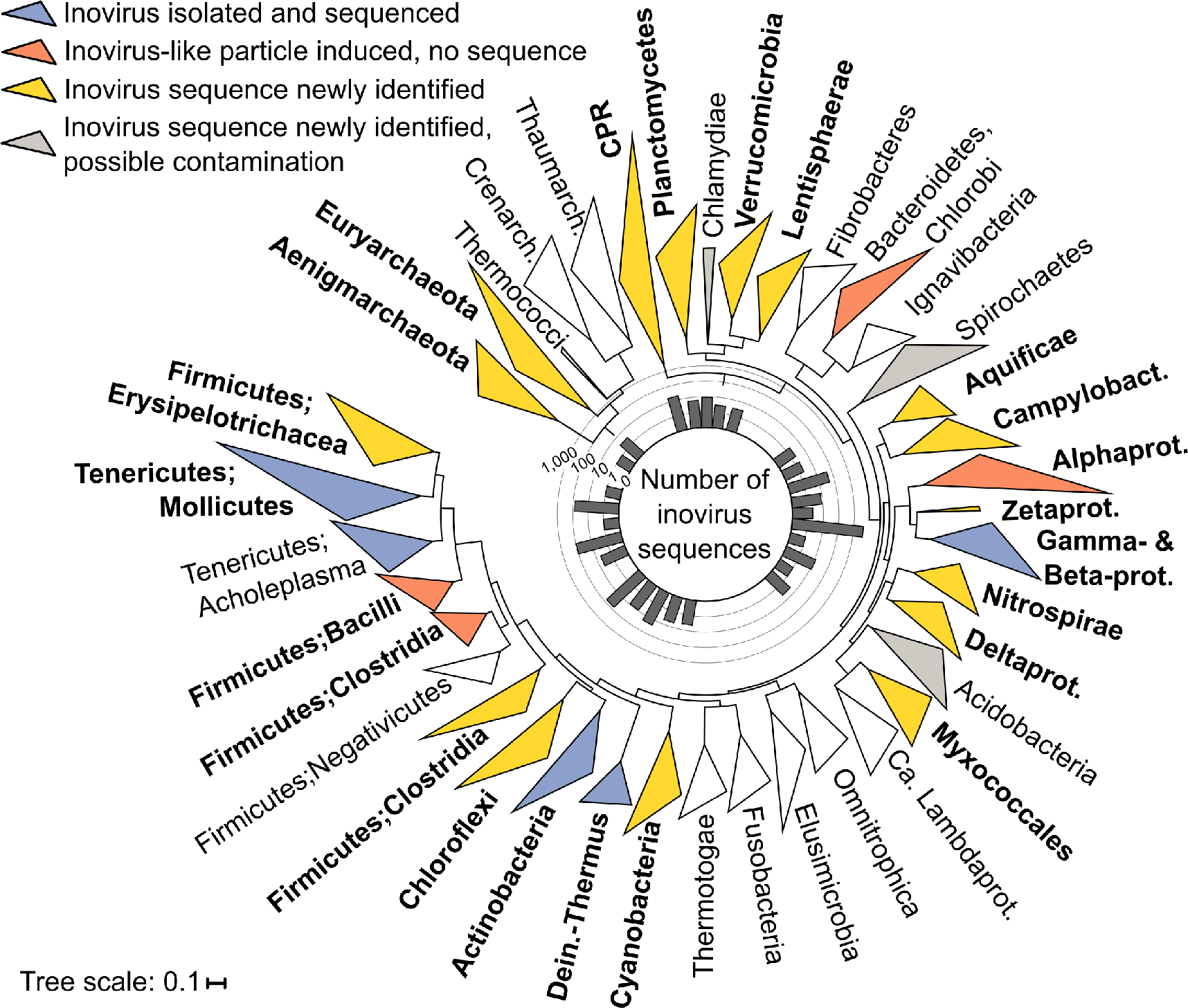
Phylum-wide distribution of inovirus detections across microbial genomes. The bacteria and archaea phylogenetic trees were computed based on 56 universal marker proteins. Monophyletic clades representing a single phylum (or class for proteobacteria) were collapsed when possible, and only clades including ≥ 30 genomes or associated with inovirus(es) are displayed. Clades for which ≥ 1 inovirus has been isolated and sequenced are colored in blue, the ones for which inovirus-like particles only have been observed are colored in orange, and clades newly associated with inoviruses are colored in yellow. Putative host clades for which inovirus detection might results from sample contamination, i.e. no clear host linkage based on on integrated prophage(s) or CRISPR spacer hit(s), are colored in gray (Table S4). Clades robustly associated with inoviruses in this study (i.e. ≥ 1 detection unlikely to result from sample contamination) are highlighted in bold. The center histogram indicates the total number of inovirus for each clade, on a log10 scale. Thaumarch.: Thaumarchaeota, Creanarch.: Crenarchaeota, CPR: Candidate Phyla Radiation, Campylobact.: Campylobacterota, Alphaprot.: Alphaproteobacteria, Zetaprot.: Zetaproteobacteria, Gamma-& Beta-prot.: Gammaproteobacteria and Betaproteobacteria, Deltaprot.: Deltaproteobacteria, Ca. Lambdaprot.: Candidatus Lambdaproteobacteria, Dein.-Thermus: Deinococcus-Thermus. See also Figure S3.

This large-scale detection of inovirus sequences in microbial genomes also enabled a comprehensive assessment of co-infection, both between different inoviruses and with other types of viruses. In the majority of cases, a single inovirus sequence was detected per genome, with multiple detections mostly found within Gammaproteobacteria, Betaproteobacteria, and *Spiroplasma* genomes (Figure S3A & B). Conversely, inovirus prophages were frequently detected along and sometimes co-localized with *Caudovirales* prophages, suggesting that these two types of phages frequently co-infect the same host cell (Figure S3C, D, & E, Supplementary Text). Overall, the broad range of bacteria and archaea infected by inoviruses combined with their propensity to co-infect a microbial cell with other viruses and their global distribution indicate that inoviruses likely play an important ecological role in all types of microbial ecosystems.

### Inoviruses sporadically transferred from bacterial to archaeal hosts

Although no archaea-infecting inoviruses have been reported so far^25^, novel hosts also included members of 2 archaeal phyla (Euryarchaeota, Aenigmarchaeota), which suggests that inoviruses infect hosts across the entire prokaryotic diversity (Figure 3). These putative archaeal proviruses encoded the full complement of genes expected in an active inovirus (Figure 4A, Supplementary Text). Using PCR, we further confirmed the presence of a circular, excised form of the complete inovirus genome for the provirus identified in the *Methanolobus profundi* MobM genome (Figure 4B, Supplementary Figure 4, Supplementary Text). This indicates that our predictions in archaeal genomes are likely genuine inoviruses.

**Figure 4.**
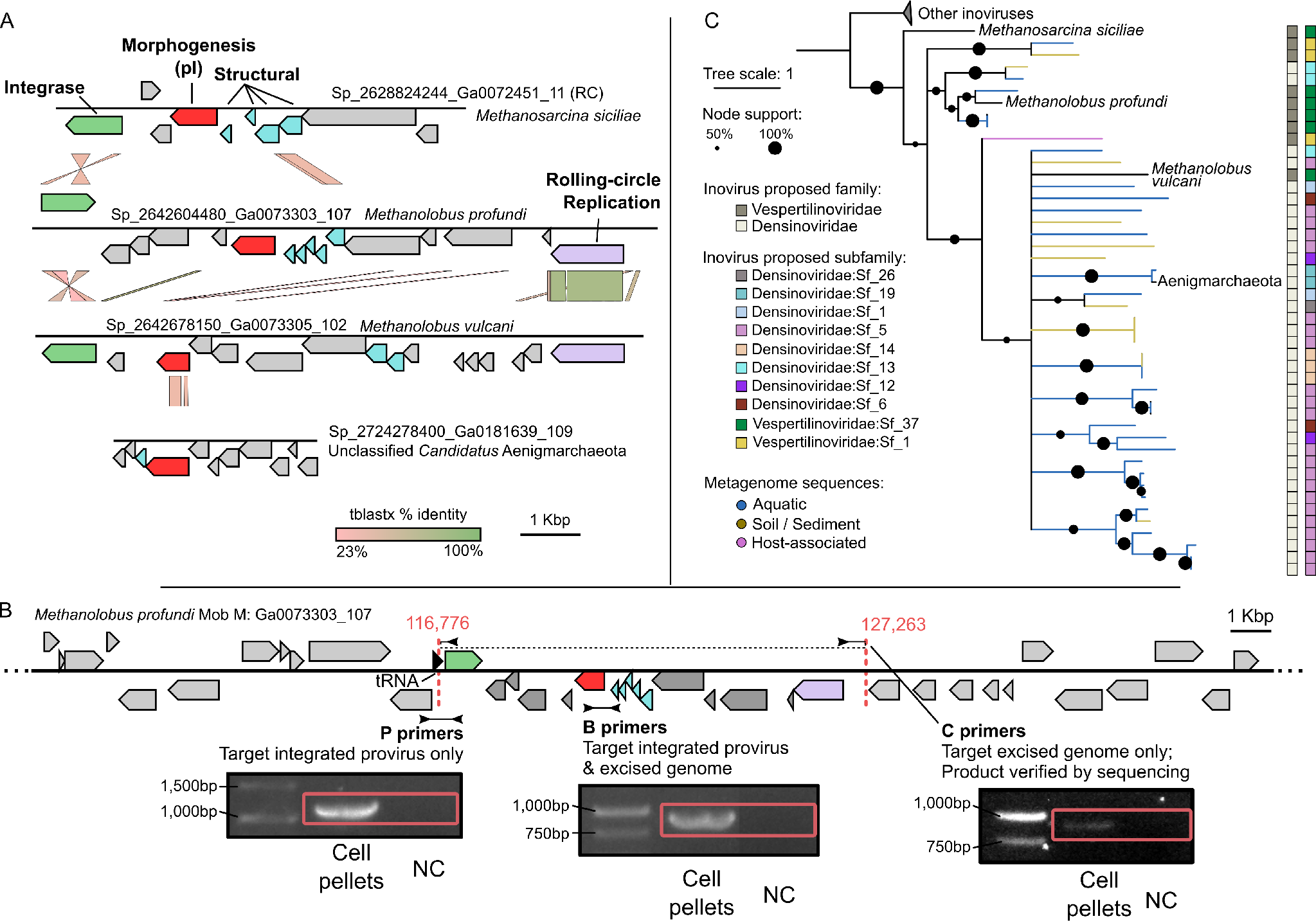
Characterization of archaea-associated inoviruses. A. Genome comparison of the 4 inovirus sequences detected in members of the *Methanosarcinaceae* family or Aenigmarchaeota candidate phylum. Genes are colored according to their functional affiliation (light gray: ORFan). RC: sequence is reverse-complemented. B. PCR validation of the predicted inovirus from the archaea host *Methanolobus profundi* MobM. Three primer pairs were designed and used to amplify across the predicted 5’ insertion site (P pirmers, left), within the predicted provirus (B primers, center) or across the junction of the predicted excised circular genome (C primers, right). Products from primers C were sequenced and aligned to the *Methanolobus profundi* MobM genome to confirm they spanned both ends of the provirus in the expected orientation and at the predicted coordinates (see Supplementary Text and Supplementary Figure S4). A red box indicates the expected product length. NC: No template control. C. Phylogenetic tree of archaea-associated inoviruses and related sequences. The tree was built from pI protein multiple alignment with IQ-tree. Nodes with support < 50% were collapsed. Branches leading to inovirus species associated to a host are colored in black, and the corresponding host is indicated on the tree. Branches leading to inovirus species assembled from metagenomes are colored by type of environment. Classification of each inovirus species in proposed families and subfamilies is indicated next to the tree (see Figure 5).

Few groups of viruses include both bacteriophages and archaeoviruses. Such evolutionary relationships between viruses infecting hosts from different domains of life might signify either descent from an ancestral virus which infected the common ancestor of bacteria and archaea, or horizontal virus transfer from one host domain to the other^25–27^. Here, the four archaea-associated inoviruses were clearly distinct from most other inoviruses and clustered only with metagenomic sequences in pI phylogeny (Figure 4C). In addition, they were classified into two different proposed families (see below) corresponding to the two host groups, reflecting clear differences in their gene content (Figure 4A & C, Supplementary Text). The high genetic diversity of these archaea-associated inoviruses, combined with the lack of similarity to bacteria-infecting species, suggest that they are not derived from a recent host switch event.

A possible scenario would involve an ancestral group of inoviruses infecting the common ancestor of archaea, as postulated for the double-jelly-roll virus lineage^27^. To be confirmed however, this hypothesis would require the detection of additional inoviruses in other archaeal clades, or an explanation as to why inoviruses were retained only in a handful of archaeal hosts. Instead, based on the current data, a more likely scenario involves ancient and rare event(s) of interdomain inovirus transfer from bacteria to archaea, including possibly to a *Methanosarcina* host for which substantive horizontal transfers of bacterial genes have already been reported^28^.

### Gene content classification reveals six distinct inovirus families

The vast increase of inovirus sequences provided a great opportunity for re-evaluation of the inovirus classification and the development of an expanded taxonomic framework for the large number of newly identified inovirus species. Similar to other bacterial viruses, especially temperate phages^29^, inovirus genomes display modular organization and are prone to recombination and horizontal gene transfers^30^ (Figure S5A). Hence, we opted to apply a bipartite network approach, in which genomes are connected to gene families, enabling a representation and clustering of the diversity based on shared gene content. A similar approach has been previously employed for the analysis of dsDNA and RNA viruses^25,31–33^. Here, this approach yielded 6 distinct groups of genomes divided into 212 sub-groups (Figure 5A, Table S3).

**Figure 5.**
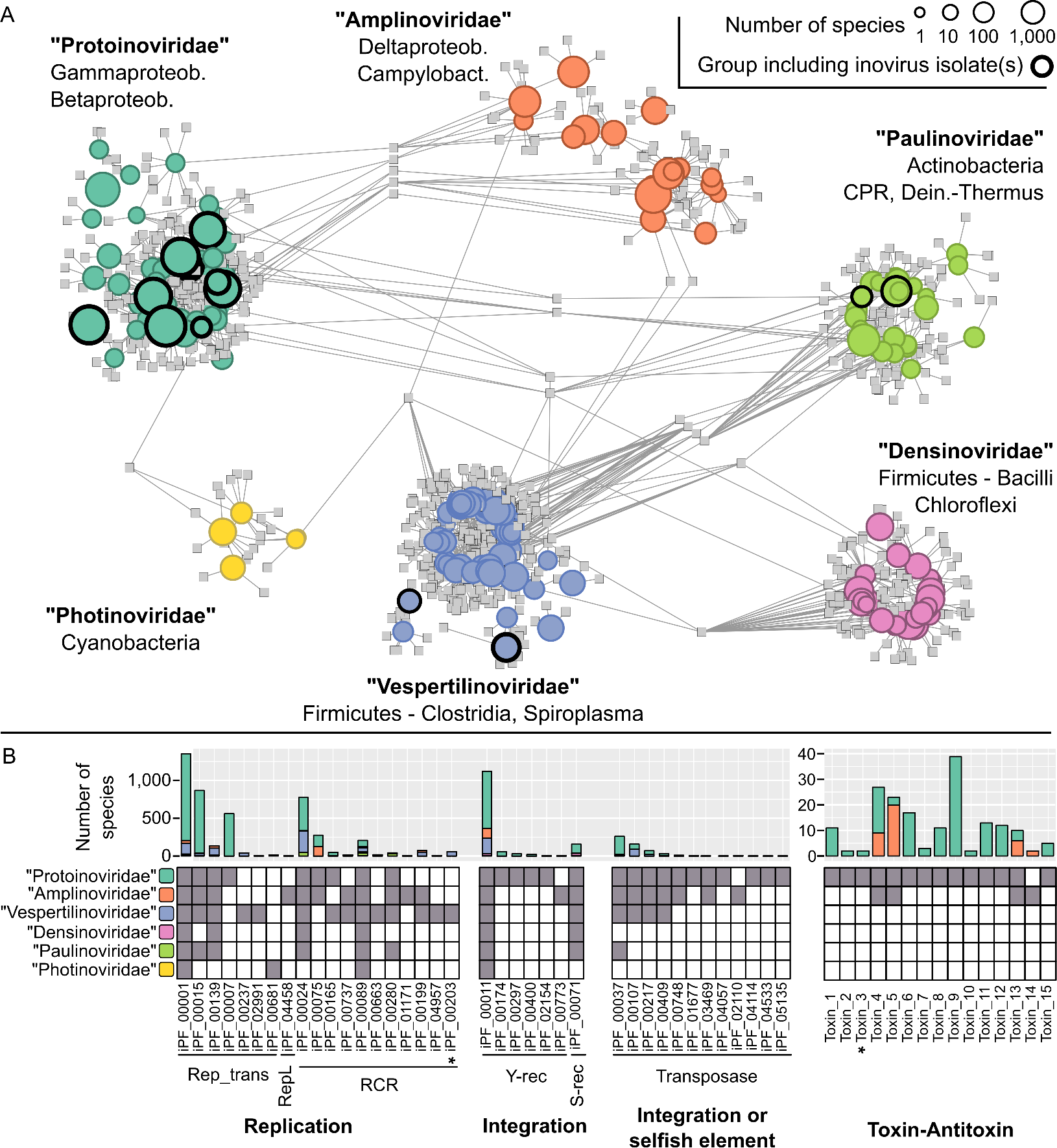
Inovirus genome sequence space and gene content. A. The bipartite network links genes represented as protein clusters (PCs) in squares, to proposed subfamilies represented as circles with a size proportional to the number of species in each candidate subfamily (log10 scale), grouped and colored by proposed family. Proposed subfamilies which include viral isolate(s) are highlighted with a black outline. Candidate subfamilies are connected to PCs when ≥ 50% of the subfamily members contained this PC, or ≥ 25% for the larger proposed subfamilies (see Methods). B. Distribution of inovirus protein families (iPFs) detected in ≥ 2 genomes, associated with genome replication, genome integration, and toxin-antitoxin systems (see Table S5). Rolling circle replication (RCR) iPFs include only the RCR endonuclease motif, with the exception of iPF_00203 (highlighted with a *), which also includes the C-terminal S3H motif typical of eukaryotic ssDNA viruses. Transposases used by selfish integrated elements are indistinguishable from transposases domesticated by viral genomes using sequence analysis only, hence these genes are gathered in a single “Integration or selfish element” category. All toxin-antitoxin pairs were predicted to be of type II, except for Toxin_3 (highlighted with a *), predicted to be type IV. RCR: rolling circle replication. Y-rec and S-rec: Tyrosine and Serine recombinase, respectively. Gammaprot.: Gammaproteobacteria, Betaprot.: Betaproteobacteria, Deltaprot.: Deltaproteobacteria, Campylobact.: Campylobacterota, CPR: Candidate Phyla Radiation, Dein.-Thermus: Deinococcus-Thermus. See also Figure S5 & S6.

A comparison of marker gene conservation between these groups and established viral taxa suggested that the former *Inoviridae* family should be reclassified as an order, provisionally divided into 6 candidate families and 212 candidate subfamilies, with few shared genes across candidate families (Figure 5A, Figure S5B, Supplementary Text). Beyond gene content, these proposed families also displayed clearly distinct host ranges as well as specific genome features, particularly in terms of genome size and coding density (Figure S5C & D). We thus propose to establish these as new candidate families named “Protoinoviridae”, “Vespertilinoviridae”, “Amplinoviridae”, “Paulinoviridae”, “Densinoviridae”, and “Photinoviridae”, based on their isolate members and characteristics (see Supplementary Text). If confirmed, and compared to currently recognized inoviruses, the new genomes reported here would increase diversity by 3 families and 198 subfamilies.

The host envelope organization appears to play an important role in the evolution of inoviruses, which is reflected in their classification: members of the “Protoinoviridae” and “Amplinoviridae” are associated with diderm hosts, i.e. Gram-negative bacteria with an outer membrane, whereas the other candidate families are associated with monoderm hosts or hosts without cell wall (Figure S5D). Conversely, no structuring by biome was observed, and all proposed families were broadly detected across multiple types of ecosystems. Hence, we propose here a classification of inovirus diversity into 6 families based on gene content with coherent host ranges and specific genome features, which strongly suggests they represent ecologically and evolutionarily meaningful units.

### Inovirus genomes encode an extensive functional repertoire

The extended catalog of inovirus genomes offers an unprecedented window into the diversity of their genes and predicted functions. Overall, 68,912 proteins were predicted and clustered into 3,439 protein families and 13,714 singletons. This is on par with the functional diversity observed in known *Caudovirales* genomes, the largest order of dsDNA viruses, for which the same number of proteins clustered into 12,285 protein families but only 8,552 singletons (see Methods). A putative function was predicted for 1,133 of the 3,439 inovirus protein families (iPFs). Most of these (> 95%) could be linked to virion structure, virion extrusion, DNA replication and integration, toxin-antitoxin systems, or transcription regulation (Table S5). A total of 51 and 47 distinct iPFs could be annotated as major and minor coat proteins, with an additional 934 iPFs identified as potentially structural based on their size and presence of a TMD (see Methods). Notably, each candidate inovirus family seemed to be associated with a specific set of structural proteins, including distinct major coat iPFs (Figure S6A). Conversely, genome replication and integration-associated iPFs were broadly shared across candidate families (Figure 5B). This confirms that replication- and integration-associated genes are among the most frequently exchanged among viral genomes and with other mobile genetic elements, especially in small ssDNA viruses^34^.

Additionally, 15 distinct sets of iPFs representing potential toxin-antitoxin (TA) pairs were identified across 181 inovirus genomes, including 10 unaffiliated iPFs which were predicted as putative antitoxins through co-occurrence with a toxin iPF (Table S5, Figure 5B, see Methods). These genes typically stabilize plasmids or prophages in host cell populations, although alternative roles in stress response and transcription regulation have been reported^35^. In addition, TA systems also often affect host cell phenotypes such as motility or biofilm formation^1^. Here, similar toxin proteins could be associated with distinct and seemingly unrelated antitoxins and vice versa, suggesting that gene shuffling and lateral transfer occur even within these tightly linked gene pairs (Figure S6B). All but one TA pairs were detected in proteobacteria-associated inoviruses, most likely because of a database bias. Thus, a number of uncharacterized iPFs across other candidate families of inoviruses may also encode novel TA systems and, more generally, include novel host manipulation mechanisms.

### Inoviruses can both leverage and restrict co-infecting viruses

Finally, we investigated potential interactions between persistently infecting inoviruses, other co-infecting viruses, and the host CRISPR-Cas immunity systems. CRISPR-Cas systems typically target bacteriophages, plasmids, and other mobile genetic elements^36^. We detected 1,150 inovirus-matching CRISPR spacers across 42 bacterial and 1 archaeal families. These spacers were associated with three types and eight subtypes of CRISPR-Cas systems indicating that inoviruses are broadly targeted by antiviral defenses (Figure 6A, Table S6, and Supplementary Text). Several host groups, most notably *Neisseria meningitidis*, displayed a particularly high ratio of inovirus-derived spacers suggesting a uniquely high level of spacer acquisition and inovirus infection (Figure 6A). This is particularly notable because inoviruses were recently suggested to increase *N. meningitidis* pathogenicity^13^, and hints at conflicting host-inovirus interactions in this specific group.

**Figure 6.**
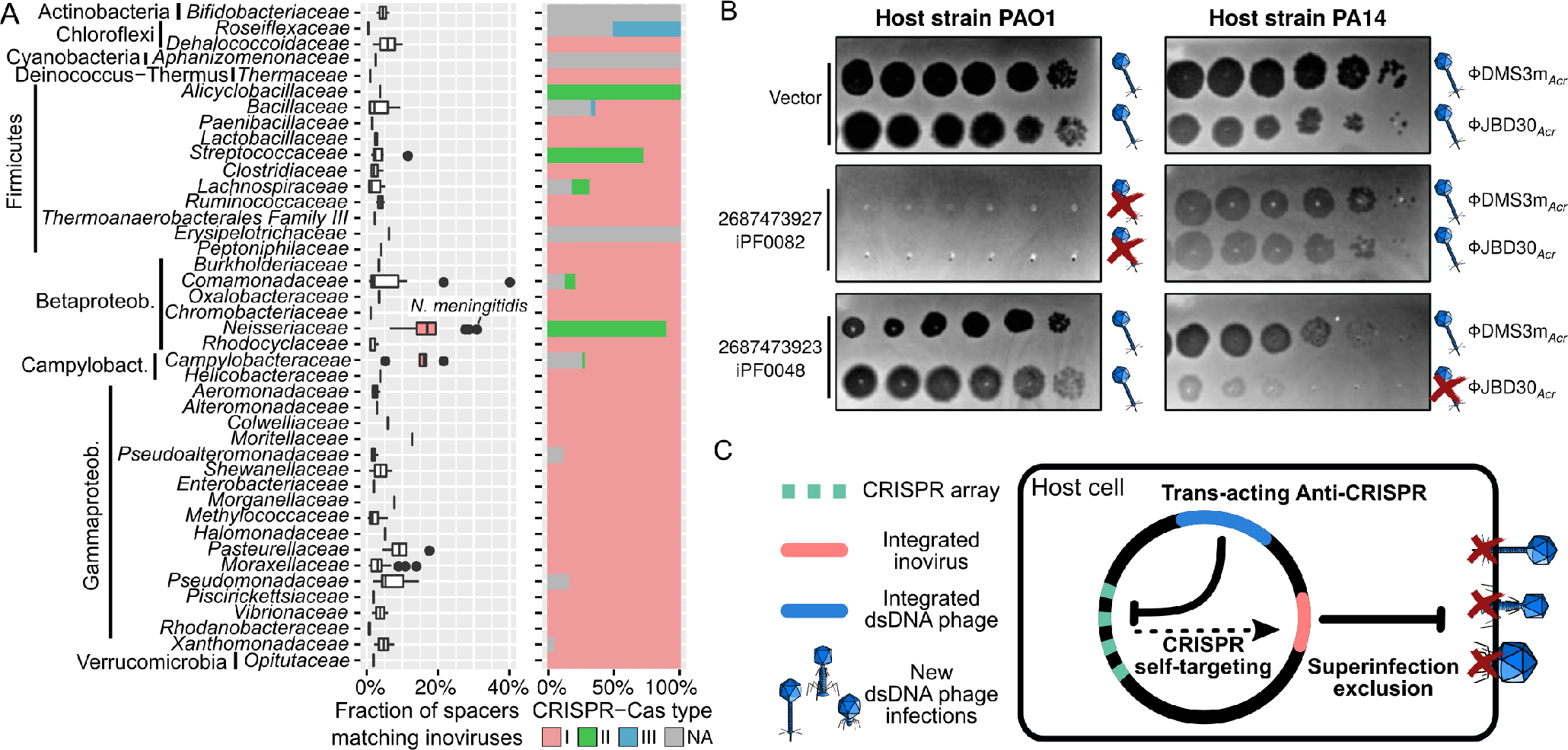
Interaction of inoviruses with CRISPR-Cas systems and co-infecting viruses. A. Proportion of the spacers matching an inovirus genome and the corresponding distribution of CRISPR-Cas systems. The proportions are calculated only on hosts with at least 1 spacer matching an inovirus sequence, with hosts grouped at the family rank (hosts unclassified at this rank were not included). The two host groups with a median inovirus-derived spacer ratio ≥ 15% are colored in red. Boxplot lower and upper hinges correspond to the first and third quartiles, whiskers extend no further than ±1.5*Inter-quartile range. B. Instances of superinfection exclusion observed when expressing individual inovirus genes in two *Pseudomonas aeruginosa* strains: PAO1 (left panel) and PA14 (right panel). From top to bottom: cells were transformed with an empty vector, one expressing gene 2687473927, or one expressing gene 2687473923. For each construct, host cells were challenged with serial dilutions (from left to right) of phages, ϕJBD30 (top row) and ϕDMS3m (bottom row). The formation of plaques (hereJBD30 (top row) and ϕJBD30 (top row) and ϕDMS3m (bottom row). The formation of plaques (hereDMS3m (bottom row). The formation of plaques (here dark circles) indicates successful infection, while the absence of plaques indicates superinfection exclusion. Interpretation of infection outcome is indicated to the right of each lane, with successful infection represented by a phage symbol, and superinfection exclusion represented by a phage symbol barred by a red cross. Results from additional superinfection exclusion experiments are presented in Figure S7. C. Schematic representation of the possible mutualistic or antagonistic interactions between inoviruses prophages (red) and co-infecting *Caudovirales* (blue). Mutualistic interactions include suppression of the CRISPR-Cas immunity, especially for integrated inoviruses targeted by the host cell CRISPR-Cas system (“self-targeting”). Antagonistic interactions primarily involve superinfection exclusion, in which a chronic inovirus infection prevents a secondary infection by an unrelated virus. Betaproteob.: Betaproteobacteria, Campylobact.: Campylobacterota.

Next, we examined instances of “self-targeting”, i.e. CRISPR spacer(s) matching an inovirus integrated in the same host genome. Among the 1,429 genomes which included both a CRISPR-Cas system and an inovirus prophage, only 45 displayed spacer match(es) to a resident prophage (Table S6), suggesting that self-targeting of these integrated elements is lethal and strongly counter-selected^37^. This was confirmed experimentally using a *Pseudomonas aeruginosa* strain PA14 harboring an integrated inovirus prophage (Pf1), for which the introduction of a plasmid carrying Pf1-targeting CRISPR spacers was lethal (Figure S7A). In the 45 cases of observed self-targeting, the corresponding CRISPR-Cas system is thus likely non-functional or inhibited via an anti-CRISPR (Acr) locus, as recently described in dsDNA phages^37^. We first evaluated 10 hypothetical proteins, hence candidate Acr proteins, from self-targeted inoviruses infecting *Pseudomonas aeruginosa*; however, none showed Acr activity (Supplementary Text, Figure S7B). Alternatively, inoviruses could leverage the Acr activity of a co-integrated virus. This hypothesis was further reinforced by the fact that 43 of the 45 self-targeted inoviruses were detected alongside co-infecting dsDNA phages, with 5 of these encoding known Acr genes (Table S6). We confirmed experimentally cross-protection by trans-acting Acr in the *Pseudomonas aeruginosa* PA14 model, and observed that co-infection with an acr-encoding dsDNA bacteriophage rescued the lethality caused by self-targeted inoviruses (Supplementary Text, Figure S7A).

While this represents an instance of beneficial co-infection for inoviruses, we also uncovered evidence of antagonistic interactions between inoviruses and dsDNA bacteriophages. Specifically, 2 of the 10 inovirus-encoded hypothetical proteins tested strongly limited infection of *Pseudomonas* cells by different bacteriophages (Figure 6B, Figure S7C, Supplementary Text). This superinfection exclusion effect was found to be host- and virus-strain dependent, which could drive intricate tripartite coevolution dynamics. These preliminary observations thus indicate that inoviruses may not only evade CRISPR-Cas immunity by leveraging the Acr activity of co-integrated phages, but also significantly influence the infection dynamics of unrelated co-infecting viruses through superinfection exclusion, with possible impacts on the host and viruses populations fitness (Figure 6C).

## Discussion

Taken together, the results presented here call for a complete re-evaluation of the diversity and role of inoviruses in nature. Collectively, inoviruses are distributed across all biomes and display an extremely broad host range spanning both prokaryotic domains of life. Comparative genomices revealed evidence of longstanding virus-host codiversification, high inovirus prevalence in several microbial groups including major pathogens, and potential interdomain transfer. Even though small (5-20kb), their genomes encode a large functional diversity shaped by frequent gene exchange with unrelated groups of viruses, plasmids and transposable elements. Some of the many uncharacterized inovirus genes likely encode novel molecular mechanisms at the interface of virus-host and virus-virus interactions such as modulators of the CRISPR-Cas systems, superinfection exclusion genes, or toxin-antitoxin modules. This expanded and restructured catalog of 5,964 distinct inovirus genomes thus provides a renewed framework for further investigation of the different impacts inoviruses have on microbial ecosystems, and exploration of their unique potential for novel biotechnological applications and manipulation of microbes.

## Methods

### Construction of an *Inoviridae* genome reference set

Genome sequences affiliated to *Inoviridae* and ≥ 2.5kb were downloaded from NCBI Genbank and RefSeq on July 14 2017^39,40^. These were clustered at 98% average nucleotide identity (ANI) to remove duplicates, and screened for cloning vectors and partial genomes (Table S1). Two of these genomes (Stenotrophomonas phage phiSMA9, NC_007189 and Ralstonia phage RSS30, NC_021862) presented unusually long section (≥ 1kb) without any predicted gene, associated with a lack of short genes that are typical of *Inoviridae.* For these, genes were predicted *de novo* using Glimmer^41^ trained on their host genomes (NC_010943 for phiSMA9, NC_003295 for RSS30) with standard genetic code. Similarly, genes for Acholeplasma phage MV-L1 (NC_001341) were predicted *de novo* using Glimmer with genetic code 4 (Mycoplasma/Spiroplasma), and trained on the host genome (NC_010163), followed by a manual curation step to integrate both RefSeq-annotated genes and these newly predicted CDS.

Protein clusters (PCs) were computed from these genomes from an all-vs-all blastp of predicted CDS (thresholds: E-value ≤ 0.001, bit score ≥ 30), and clustered with InfoMap^32^. Sequences from these protein clusters were then aligned with Muscle ^42^, transformed into an HMM profile and compared to each other using HHSearch^43^ (cutoffs: probability ≥ 90% and coverage ≥ 50%, or probability ≥ 99%, coverage ≥ 20%, and hit length ≥ 100). The larger clusters generated through this second step are designated here as inovirus protein families (iPFs). Only 10 PCs were clustered into larger iPFs, but these were consistent with the functional annotation of these proteins. For instance, 1 iPF combined 2 PCs both composed of replication initiation proteins.

Marker genes were identified from a bipartite network linking *Inoviridae* genomes to iPFs (Figure S1A). Only the genes coding for the morphogenesis (pI) protein represented good candidates for a universally conserved gene across all members of the *Inoviridae*, and HMM profiles were built for the 3 pI iPFs. To optimize these profiles, sequences were first clustered at 90% AAI with cd-hit^44^, then aligned with Muscle ^42^ and the profile generated with hmmbuild^45^.

These reference genomes were also used to evaluate the detection of the *Inoviridae* structural proteins based on protein features beyond sequence similarity (see Supplementary Text). Here, signal peptides were predicted using SignalP in both gram positive and gram negative modes^46^, and transmembrane domains were identified with TMHMM^47^.

### Search for inovirus in microbial genomes and metagenomes

Proteins predicted from 56,868 microbial genomes publicly available in IMG as of October 2017 (Table S2) were compared to the reference morphogenesis (pI) proteins with hmmsearch^45^ (hmmer.org, score ≥ 30 E-value ≤ 0.001) for the pI-like iPFs and blastp^48^ (bit score ≥ 50) for the singleton pI protein (Acholeplasma phage MV-L1). These included 54,405 bacterial genomes, 1,304 archaeal genomes, and 1,149 plasmid sequences. A total of 6,819 hits were detected, from which 795 corresponded to complete inovirus genomes. These included 213 circular contigs, i.e. likely complete genomes, and 582 integrated prophages with canonical attachment (att) sites, i.e. direct repeats ≥ 10bp in a tRNA or outside of an integrase gene. All sequences were manually inspected to verify that these were plausible inovirus genomes (see Supplementary Text). The predicted pI proteins from the curated genomes were then added to the references to generate new improved HMM models. Using these improved models, an additional set of 639 putative pI proteins was identified. New models were built from these proteins and used in a third round of searches, which did not yield any additional genuine inovirus sequence after manual inspection.

An automatic classifier was trained on this extended inovirus genome catalog, i.e. the reference genomes and the 795 manually curated genomes, to detect putative inovirus fragments around pI-like genes, based on 10 distinctive features of inovirus genomes (Figure S1B, see Supplementary Text). These 795 manually curated genomes were identified from 17 host phyla (or class for Proteobacteria), and were later classified into 5 proposed families and 245 proposed subfamilies (see below “Gene-content based clustering of inovirus genomes”). Three types of classifiers were tested: Random forest (function randomForest from R package randomForest^49^), Random forest with conditional inference (function cforest from R package party^50^), and a Generalized linear model with lasso regularization (function glmnet from R package glmnet^51^). The efficiency of classifiers was evaluated via a 10-fold cross-validation, and results were visualized as a ROC curve generated with ggplot2^52,53^.

Based on the inflection point observed on the ROC curves, the random forest classifier was selected as the optimal method since it provided the highest true positive rate (> 92%) for false positive rates < 1 % (Figure S1 C). This model was then used to classify all putative inovirus fragments that had not been identified as complete genomes previously, using a sliding window approach (up to 30 genes around the putative pI protein), and looking for the fragment with the maximum score in the random forest model (if > 0.9). For the predicted integrated prophages, putative non-canonical att sites were next searched as direct repeats (10bp or longer) around the fragment. Overall, 3,908 additional putative inovirus sequences were detected, including 738 prophages flanked by direct repeats.

A similar approach was used to search for inovirus sequences in 6,412 metagenome assemblies (Table S2). Predicted proteins were compared to the 4 HMM profiles as well as to the Acholeplasma phage MV-L1 singleton sequence, which led to 27,037 putative pI proteins using the same thresholds as for isolate genomes. The final dataset of inovirus sequences predicted from these metagenome assemblies consisted of 6,094 sequences including 922 circular contigs, 44 prophages with canonical att sites (direct repeats of 10bp or longer in a tRNA or next to an integrase), and 994 prophages with non-canonical att sites (direct repeats of 10bp or longer).

### Clustering of inovirus genomes in putative species

Next, we sought to cluster these putative inovirus genomes along with the previously collected reference genomes in order to remove duplicated sequences and to select only one representative per species. This clustering was conducted according to the latest guidelines submitted to the ICTV for *Inoviridae*, i.e. “95% DNA sequence identity as the criterion for demarcation of species”^54^, and included our 10,295 sequences alongside the 56 reference genomes. Notably, however, predictions spanning multiple tandemly integrated inovirus prophages had to be processed separately; otherwise, they could lead to clusters gathering multiple species. To detect these cases of tandem insertions, we searched for and clustered separately all predictions with multiple pI proteins, as this gene is expected to be present in single copy in inoviruses (n=800 sequences).

All non-tandem sequences were first clustered incrementally with priority given to complete genomes over partial genomes as well as fragments identified in microbial genomes over fragments from metagenomes. First, circular contigs and prophages with canonical att sites identified in a microbial genome were clustered, and all other fragments were affiliated to these seed sequences. Next, unaffiliated fragments detected in microbial genomes and with non-canonical att sites (i.e. simple direct repeat) were clustered together, and other fragments were affiliated to this second set of seed sequences. Finally, the remaining unaffiliated sequences detected in microbial genomes were clustered together. This allowed us to use the more “certain” predictions (i.e. circular sequences and prophages with identified att sites) preferentially as seeds of putative species.

A similar approach was used to cluster sequences identified from metagenomes, as well as to separately cluster putative tandem fragments, i.e. those including multiple pI proteins. All the clustering and affiliation was done with a threshold of 95% ANI on 100% of alignment fraction (according to the ICTV guidelines), with sequence similarity computed using mummer^55^. Accumulation curves were calculated for 100 random ordering of input sequences using a custom perl script, and plotted with ggplot2^52,53^.

### Clustering of predicted proteins from non-redundant inovirus sequences

Predicted proteins from the representative genome of each putative species were next clustered using the same approach as for the reference genomes. A clustering into protein clusters (PCs) was first achieved through an all-vs-all blastp using hits with ≤ 0.001 e-value and ≥ 50 bit score or ≥ 30 bit score if both proteins are ≤ 70 aa. HMM profiles were constructed for the 5,142 PCs, and these were compared all-vs-all using HHSearch, keeping hits with ≥ 90% probability and ≥ 50% coverage or ≥ 99% probability, ≥ 20% coverage, and hit length ≥ 100. This resulted in 4,008 protein families (iPFs).

The PCs were subsequently used for taxonomic classification of the inovirus sequences (see below), while iPFs were primarily used for functional affiliation. iPF functions were predicted based on the affiliation of iPF members against PFAM v30 (score ≥ 30), as well as manual inspection of individual iPFs using HHPred^56^.

PCs containing pI-like protein(s) were also further evaluated to identify potential false-positives stemming from a related ATPase encoded by another type of virus or mobile genetic element (see Supplementary Text). The criteria used to determine a genuine inovirus pI-like PCs were: the PCs members closest known functional domain was Zot (based on the hmmsearch against PFAM), the proteins contained 1 or 2 TMD (either N-terminal or C-terminal), at least half of the sequences encoding this PC also include other genes expected in an inovirus sequence such as replication initiation proteins, and no significant similarity could be identified to any other type of ATPase using HHpred^56^.

### Gene-content based clustering of inovirus genomes

A bipartite network was built in which genomes and PCs (as nodes) are connected by an edge when a predicted protein from the genome is a member of the PC. This network was then used to classify inovirus sequences as done previously for dsDNA viruses^31^. PCs were used instead of iPFs as they offer a higher resolution. Sequences with two pI proteins (i.e. tandem prophages) were excluded from this network-based classification as these could lead to improper connections between unrelated genomes. Singleton proteins were also excluded, and only PCs with at least 2 members were used to build the network. This network had a very low density (0.05%) reflecting the fact that most PCs were restricted to a minor fraction of the genomes. Nevertheless, this type of network can still be organized into meaningful groups through information theoretic approaches: here, sequence clusters were obtained through InfoMap, with default parameters and a 2-level clustering, i.e. genomes can be associated with a group and a subgroup.

A summarized representation of the network was generated by displaying each sub-group (level 2) as a node with a size proportional to the number of species in the sub-group, and drawing an edge to a PC if > 50% of the sub-group sequences encode this PC, except for the larger group (“Protoinoviridae”:Subfamily_1) where connections are drawn for PCs found in > 25% of the sequences. The network was then visualized using Cytoscape^57^, with nodes from the same group (level 1) first gathered manually, and nodes allotment within group automatically generated using Prefuse directed layout (default spring length 200).

To evaluate the taxonomic rank to which these groups and sub-groups would correspond, we calculated pairwise amino acid identity percentage (AAI) of pI proteins for genomes (i) between groups and (ii) within groups but between sub-groups, using SDT^58^. These were then compared to pairwise AAI calculated with the same approach for established viral groups, namely *Caudovirales* order using the Terminase large subunit (TerL) as a marker protein, *Microviridae* using the major capsid protein (VP1) as a marker protein, and *Circoviridae* using the replication initiation protein (Rep) as a marker protein (see Supplementary Text).

### Distribution of inovirus sequences by host and biome

The distribution of hosts for inovirus sequences was based on detections in IMG draft and complete genomes, i.e. excluding all metagenome-derived detections but including detections in metagenome-assembled genomes (published draft genomes assembled from metagenomes). Host taxonomic classification was extracted from the IMG database. For visualization purposes, a set of 56 universal single copy marker proteins^59,60^ was used to build phylogenetic trees for bacteria and archaea based on all available microbial genomes in IMG^22^ (genomes downloaded 27 October 2017) and about 8,000 metagenome assembled genomes from the Genome Taxonomy Database^61^ (downloaded 18 October 2017). Marker proteins were identified with hmmsearch (version 3.1b2, hmmer.org) using a specific hmm for each of the markers. Genomes lacking a substantial proportion of marker proteins (> 28) or which had additional copies of > 3 single-copy markers were removed from the data set.

To reduce redundancy and to enable a representative taxon sampling, DNA directed RNA polymerase beta subunit 160kD (COG0086) was identified using hmmsearch (hmmer 3.1b2) and the HMM of COG0086^62^. Protein hits were then extracted and clustered with cd-hit^44^ at 65% sequence similarity, resulting in 99 archaeal and 837 bacterial clusters. Genomes with the greatest number of different marker proteins were selected as cluster-representatives. For every marker protein, alignments were built with MAFFT^63^ (v7.294b) and subsequently trimmed with BMGE (v1.12) using BLOSUM30^64^. Single protein alignments were then concatenated resulting in an alignment of 11,220 sites for the archaea and 16,562 sites for the bacteria. Maximum likelihood phylogenies were inferred with FastTree2 (v2.1.9 SSE3, OpenMP)^65^ using the options: -spr 4 -mlacc 2 -slownni -lg.

A distribution of inovirus sequences across biomes was obtained by compiling ecosystems and sampling location of all metagenomes where at least one inovirus sequence was detected. This information was extracted from the GOLD database^66^, and the map generated using the BaseMap functions from the matplotlib python library^67^.

### Estimation of inovirus prevalence and co-infection patterns

Prevalence and co-infection patterns were evaluated from the set of sequences identified in complete and draft microbial genomes from the IMG database, i.e. excluding detections from metagenome assemblies. To control for the presence of near-identical genomes in the database, prevalence and co-infection frequencies were calculated after clustering host genomes based on pairwise ANI (cutoffs: 95% nucleotide identity on 95% alignment fraction). Prevalence was calculated at the host genus rank as the number of genomes with ≥ 1 inovirus sequence detected. Co-occurrence of inoviruses was evaluated based on the detections of distinct species in single host genomes. Finally, we evaluated the rate of bacteria and archaea co-infected by an inovirus and a member of the *Caudovirales* order, the group of dsDNA viruses including most of characterized bacteriophages (both lytic and temperate) as well as several archaeoviruses. To identify *Caudovirales* infections, we used the gene coding for the Terminase large subunit as a marker gene, and searched the same genomes from the IMG database for hits to the PFAM domains Terminase_1, Terminase_3, Terminase_6, and Terminase_GpA (hmmsearch, score ≥ 30).

### Phylogenetic trees of inovirus sequences

Phylogenies of inovirus sequences were based on multiple alignment of pI protein sequences. To obtain informative multiple alignments, an all-vs-all blastp^48^ of all pI proteins was computed and used to identify the nearest neighbors of sequences of interests. For sequences detected in archaeal genomes, an additional 10 most closely related sequences with e-value ≤ 10^-3^, bit score ≥50, and blast hit covering ≥ 50% of the query sequence were recruited for each archaea-associated sequence to help populate the tree. A similar approach was used for the tree based on the Integrase genes from archaea-associated inoviruses: the protein sequences for the three integrase genes were compared to the NCBI nr database with blastp^48^ (bit score ≥ 50, e-value ≤ 0.001) to gather their closest neighbors across archaeal and bacterial genomes.

Resulting datasets were first filtered for partial sequences as follows: the average sequence length was calculated excluding top and bottom 10%, and all sequences shorter than half of this average were excluded. These protein sequences were next aligned with Muscle (v3.8.1551)^42^, automatically trimmed with trimAL (v1.4.rev15)^68^ (option gappyout), and trees were constructed using IQ-Tree (v1.5.5) with an automatic detection of optimal model^69^, and displayed using iToL^70^. The optimal substitution model, selected based on the Bayesian Information Criterion, was VT+F+R5 for the the pI phylogeny of archaeal inoviruses, and LG+R4 for the integrase phylogeny of archaeal inoviruses. Annotated trees are available at http://itol.embl.de/shared/Siroux (project “Inovirus”).

### Functional affiliation of inovirus protein families (iPFs)

An automatic functional affiliation of all iPFs was generated by compiling the annotation of all members based on a comparison to PFAM (data extracted from IMG). To refine these annotations for functions of interest, namely Replication initiation proteins, Integration proteins, DNA methylases, and Toxin-antitoxin systems, individual iPF alignments were submitted to the HHPred website^56^, and the alignments were visually inspected for conserved residues and/or motifs (Table S5, motifs extracted from refs.^71,72^, and the PFAM database v30^73^).

To identify toxin-antitoxin protein partners, all inovirus sequences were screened for co-occurring genes including an iPF annotated as toxin and/or antitoxin, and the list of putative pair was next manually curated (Table S5). This enabled the identification of putative novel antitoxin proteins detected as conserved uncharacterized iPF frequently observed next to a predicted toxin iPF.

Finally, novel structural proteins and DNA-interacting proteins were specifically searched for. Putative structural proteins were predicted as described above for the isolate reference genomes, i.e. as sequences of 30 to 90 amino acids, after in silico removal of signal peptide, if detected, and displaying 1 or 2 TMD. For the most abundant iPFs predicted as major coat proteins, secondary structure was predicted with Phyre2^74^. For DNA-interacting proteins, PFAM annotations were screened for HTH, RHH, Zn-binding, and Zn-ribbon domains. In addition, HHsearch was used to compare the iPFs to 3 conserved HTH domains from the SMART database^75^: Bac_DnaA_C, HTH_DTXR, and HTH_XRE (probability ≥ 90).

### CRISPR spacer matches and CRISPR-Cas systems identification

All inovirus sequences were compared to the IMG CRISPR spacer database with blastn, using options adapted for short sequences (“-task blastn-short -evalue 1 -word_size 7 -gapopen 10 -gapextend 2 -penalty -1 -dust no). Only cases with 0 or 1 mismatch were further considered. Next, the genome context of these spacers was explored to identify the ones with a clear associated CRISPR-Cas system, and affiliate these systems to the different types described. Only spacers for which a Cas gene could be identified in a region of ±10kb were retained. The CRISPR-Cas system affiliation was based on the set of Cas genes identified around the spacer and performed following the guidelines from ref.^76^.

For host genomes with a self-targeting spacer, additional (i.e. non-inovirus) prophages were detected using VirSorter^19^. The number of distinct prophages was also estimated using the detection of large terminase subunits (hmmsearch against PFAM database, score ≥ 30). Putative Anti-CRISPR (Acr) and associated (Aca) proteins were first detected through similarity to previously described Acr systems^37^ (blastp, e-value ≤ 0.001 and score ≥ 50). Putative novel Acr and Aca proteins were identified by searching for HTH-domain-containing proteins identified based on HTH domains in the SMART database (see above) in inovirus sequences displaying a match to a CRISPR spacer extracted from the same host genome.

### Microscopy and PCR investigation of predicted provirus in *Methanolobus profundi* MobM

*Methanolobus profundi* strain MobM cells were grown in anaerobic DSMZ medium 479 at 37°C with 5mM methanol added as a methanogenic substrate instead of trimethylamine^38^. After 35 hours of growth, anaerobic mitomycin C was added to the culture at a final concentration of 1.0 μg/mL tog/mL to induce the provirus. Samples were collected before and 4 hours after induction and were filtered with 0.22 μg/mL tom pore size polyethersulfone (PES) filters (Millipore, Fisher Scientific) to obtain a “cellular” (≥ 0.22 μg/mL tom) and a “viral” (< 0.22 μm) fraction.

The 4 types of samples (with or without induction, cellular and viral fractions) were prepared and imaged at the Molecular and Cellular Imaging Center, Ohio State University, Wooster OH. An equal volume of 2x fixative (6% glutaraldehyde, 2% paraformaldehyde in 0.1 M potassium phosphate buffer pH 7.2) was added directly to the culture post-induction. 30 μg/mL tol of medium was applied to a formovar and carbon coated copper grid for 5 minutes, blotted and then stained with 2% uranyl acetate for 1 minute. Samples were examined with a Hitachi H7500 electron microscope and imaged with the SIA-L12C (16 megapixels) digital camera.

PCR reactions were initially run for induced and non-induced samples on both size fractions with three pairs of primers: one internal to the predicted provirus (B primers), one spanning the insertion site (P primers), and one spanning the junction of the predicted excised circular genome (C primers). The reactions were conducted for 35 cycles with denaturation, annealing, and extension cycles of 0.5, 0.5, and 1.0 minutes at 95.0, 52.0, and 72.0°C, respectively. For primers C, a number of nonspecific amplification products were obtained with these conditions, and another set of PCR reaction were conducted with higher annealing temperatures of 56.5°C and 57.5°C, both in triplicates. The PCR product was then cleaned to remove polymerase, free dNTPs and primers (Zymo Research) and subsequently used as templates for Sanger sequencing. The resulting chromatograms were analyzed using the R^53^ packages sangerseqR^77^, sangeranalyseR^78^, and readr^79^. The extracted primary sequences were aligned to the MobM genome using blastn^48^ and muscle^42^, and the alignment visualized with Jalview^80^.

### Experimental characterization of hypothetical proteins from self-targeted *Pseudomonas* inoviruses

Hypothetical proteins predicted on inovirus prophages which were (i) found in *Pseudomonas* genomes, (ii) predicted to be targeted by at least 1 CRISPR spacer from the same genome, and (iii) for which no anti-CRISPR locus could be identified anywhere else in the same genome, were selected for further functional characterization. The 10 candidate genes were first codon-optimized for expression in *Pseudomonas* using an empirically derived codon usage table. Codon optimization and vendor defined synthesis constraints removal were performed using BOOST^81^. Synthetic DNA were obtained from Thermo Fisher Scientific and cloned in between the SacI and PstI sites of an *Escherichia*-*Pseudomonas* broad-host-range expression vector, pHERD30T^82^. All gene constructs were sequence-verified before testing.

*Pseudomonas aeruginosa* strains (PAO1::pLac I-C CRISPR-Cas, PA14, and 4386) were cultured on lysogeny broth (LB) agar or liquid media at 37 °C. The pHERD30T plasmids were electroporated into *Pseudomonas aeruginosa* strains, and LB was supplemented with 50 μg/mL gentamicin to maintain the pHERD30T plasmid. Phages DMS3m, JBD30, D3, 14-1, Luz7, and KMV were amplified on PAO1 and phage JBD44a was amplified on PA14. All phages were stored in SM buffer at 4 °C in the presence of chloroform.

For phage titering, a bacterial lawn was first generated by spreading 6 mL of top agar seeded with 200 μl of host bacteria on a LB agar plate supplemented with 10 mM MgSO4, 50 μg/mL gentamicin, and 0.1% arabinose. The I-C Cas genes in strain PAO1 were induced with 1mM Isopropyl ß-D-1-thiogalactopyranoside (IPTG). 3 μl of phage serially diluted in SM buffer was then spotted onto the lawn, and incubated at 37 ºC for 16 hours. Growth rates were similar between cells transformed with an empty vector and cells transformed with a vector including a candidate gene, except for the two cases where no growth was observed after transformation (see Supplementary Text).

### Experimental confirmation of self-targeting lethality and trans-acting Acr activity from co-infecting phage in a *Pseudomonas aeruginosa* model

The impact of CRISPR targeting of an integrated inovirus prophage was assessed in a *Pseudomonas aeruginosa* strain PA14 which naturally encodes an intact Pf1 inovirus prophage, and for which both natural CRISPR arrays were deleted (strain PA14 ∆CRISPR1/∆CRISPR2 [Pf1]). Host cells were transformed with plasmids encoding CRISPR spacers either targeting Pf1 coat gene or without a target in the host genome. To generate these plasmids, complementary single stranded oligos (IDT) were annealed and ligated into a linearized derivative of shuttle vector pHERD30T bearing I-F direct repeats in the multiple cloning site downstream of the pBAD promoter. PA14 lysogens were electroporated with 100ng of plasmid DNA, allowed to recover for 1 hour in LB at 37 °C, and plated on LB agar plates supplemented with 50 μg/mL tog/mL gentamicin and 0.1% arabinose. Colonies were enumerated after growth for 14 hours at 37 °C. Transformation efficiency (TE) was calculated as colonies/ μg/mL tog DNA, and % TE was calculated by normalizing the TE of the crRNA-expressing plasmids to the TE of an empty vector.

To evaluate the impact of an Acr locus from a co-infecting prophage on self-targeted inoviruses, strain PA14 ∆CRISPR1/∆CRISPR2 [Pf1] was lysogenized with phage DMS3m*acrIF1* by streaking out cells from a solid plate infection and screening for colonies resistant to superinfection by DMS3m*acrIF1*. Lysogeny was confirmed by prophage induction. The same plasmid transformation approach was then used to assess the impact of inovirus self-targeting on host cell viability.

### Quantification and statistical analysis

Sequence similarity searches were conducted with thresholds of 0.001 on E-value and 30 or 50 on bit score, the former being used mainly for short proteins. The different classifiers (Random Forest, Conditional Random Forest, and Generalized Linear Model) used to identify inovirus sequences were evaluated using a 10-fold cross-validation approach. For all boxplots, lower and upper hinges correspond to the first and third quartiles, and whiskers extend no further than ±1.5*Inter-quartile range.

### Data and software availability

Gb_files_inoviruses.zip: GenBank files of all representative genomes for each inovirus species.

Ref_PCs_inoviruses.zip: Protein clusters from the references (raw fasta, alignment fasta, hmm profile).

iPFs_inoviruses.zip: Protein families from extended inovirus dataset (raw fasta, alignment fasta, hmm profile).

MobM_C_primer_amplicon.fasta: Multiple sequence alignment of the C primer products with *Methanolobus* MobM genome (NZ_FOUJ01000007) confirming that C primer products span the junction of the excised genome.

These files are currently available for review at https://tinyurl.com/ybpoq4hw, and will be made available through a custom JGI genome portal upon publication, similar as was done e.g. for the PhyloTag project https://genome.jgi.doe.gov/portal/PhyloTag/PhyloTag.home.html. The set of novel scripts and models used to detect novel inovirus sequences are available at https://github.com/simroux/Inovirus/tree/master/Inovirus_detector.

## Acknowledgments

The MobM strain was provided by D.J. Ferguson, Miami University, Oxford, OH. Its genome was sequenced and assembled by the US Department of Energy Joint Genome Institute through a Community Science Program initiative to K.C.W. (CSP #1777). Tea Meulia at the Molecular and Cellular Imaging Center, Ohio State University, Wooster OH performed the TEM of MobM samples.

## Funding

This work was conducted by the US Department of Energy Joint Genome Institute, a DOE Office of Science User Facility, under Contract No. DE-AC02–05CH11231 and used resources of the National Energy Research Scientific Computing Center, which is supported by the Office of Science of the US Department of Energy under Contract No. DE-AC02–05CH11231. R.A.D. and K.C.W. were partially supported by funding from the National Sciences Foundation Dimensions of Biodiversity (Award 1342701). M.K. was supported by l’Agence Nationale de la Recherche (France) project ENVIRA. The Bondy-Denomy lab (A.L.B. and J.B.-D.) is supported by the UCSF Program for Breakthrough in Biomedical Research, funded in part by the Sandler Foundation, the NIH Office of the Director (DP5-OD021344), and NIGMS (R01GM127489).

## Author contributions

All authors participated in writing and reviewing the manuscript. S.R., M.K., and E.A.E.-F. conceived the study. S.R. and M.K. performed the data and metadata curation. S.R. developed the novel computational tools. S.N. and F.S. contributed additional computational analyses. R.A.D., J.-F. C., and A.L.B. performed the experiments.

## Declaration of Interests

The authors declare no competing interests

## References

1. Rakonjac, J., Bennett, N. J., Spagnuolo, J., Gagic, D. & Russel, M. Filamentous bacteriophage: biology, phage display and nanotechnology applications. Curr. Issues Mol. Biol. 13, 51–76 (2011).

2. Fauquet, C. M. The diversity of single stranded DNA viruses. Biodiversity 7, 38–44 (2006).

3. Marvin, D. A., Symmons, M. F. & Straus, S. K. Structure and assembly of filamentous bacteriophages. Prog. Biophys. Mol. Biol. 114, 80–122 (2014).

4. Mai-Prochnow, A. et al. ‘Big things in small packages: The genetics of filamentous phage and effects on fitness of their host’. FEMS Microbiol. Rev. 39, 465–487 (2015).

5. Bradbury, A. R. M. & Marks, J. D. Antibodies from phage antibody libraries. J. Immunol. Methods 290, 29–49 (2004).

6. Nam, K. T. et al. Stamped microbattery electrodes based on self-assembled M13 viruses. Proc. Natl. Acad. Sci. 105, 17227–17231 (2008).

7. Ju, Z. & Sun, W. Drug delivery vectors based on filamentous bacteriophages and phage-mimetic nanoparticles. Drug Deliv. 24, 1898–1908 (2017).

8. Henry, K. A., Arbabi-Ghahroudi, M. & Scott, J. K. Beyond phage display: Non-traditional applications of the filamentous bacteriophage as a vaccine carrier, therapeutic biologic, and bioconjugation scaffold. Front. Microbiol. 6, 1–18 (2015).

9. Ilyina, T. S. Filamentous bacteriophages and their role in the virulence and evolution of pathogenic bacteria. Mol. Genet. Microbiol. Virol. 30, 1–9 (2015).

10. Shapiro, J. W. & Turner, P. E. Evolution of mutualism from parasitism in experimental virus populations. Evolution (N. Y). 72, 707–712 (2018).

11. Waldor, M. K. & Mekalanos, J. J. Lysogenic conversion by a filamentous phage encoding cholera toxin. Science 272, 1910–1914 (1996).

12. Faruque, S. M. & Mekalanos, J. J. Pathogenicity islands and phages in Vibrio cholerae evolution. Trends Microbiol. 11, 505–510 (2003).

13. Bille, E. et al. A virulence-associated filamentous bacteriophage of Neisseria meningitidis increases host-cell colonisation. PLoS Pathog. 13, 1–23 (2017).

14. Rice, S. A. et al. The biofilm life cycle and virulence of Pseudomonas aeruginosa are dependent on a filamentous prophage. ISME J. 3, 271–282 (2009).

15. Rakonjac, J. Filamentous Bacteriophages: Biology and Applications. eLS (2012). doi:10.1002/9780470015902.a0000777

16. Varani, A. M., Monteiro-Vitorello, C. B., Nakaya, H. I. & Van Sluys, M.-A. The Role of Prophage in Plant-Pathogenic Bacteria. Annu. Rev. Phytopathol. 51, 429–451 (2013).

17. Páez-Espino, D. et al. Uncovering Earth’s virome. Nature 536, 425–430 (2016).

18. Páez-Espino, D., Pavlopoulos, G. A., Ivanova, N. N. & Kyrpides, N. C. Nontargeted virus sequence discovery pipeline and virus clustering for metagenomic data. Nat. Protoc. 12, 1673–1682 (2017).

19. Roux, S., Enault, F., Hurwitz, B. L. & Sullivan, M. B. VirSorter: mining viral signal from microbial genomic data. PeerJ 3, e985 (2015).

20. Brum, J. R. & Sullivan, M. B. Rising to the challenge: accelerated pace of discovery transforms marine virology. Nat. Rev. Microbiol. 13, 1–13 (2015).

21. Vega Thurber, R. V et al. Laboratory procedures to generate viral metagenomes. Nat. Protoc. 4, 470–483 (2009).

22. Chen, I. M. A. et al. IMG/M: Integrated genome and metagenome comparative data analysis system. Nucleic Acids Res. 45, D507–D516 (2017).

23. Kimura, M., Wang, G., Nakayama, N. & Asakawa, S. in Biocommunication in Soil Microorganisms. (ed. Witzany, G.) 189–213 (Springer Berlin Heidelberg, 2011).

24. Kim, A. Y. & Blaschek, H. P. Isolation and characterization of a filamentous virus-like particle from Clostridium acetobutylicum Ncib-6444. J. Bacteriol. 173, 530–535 (1991).

25. Iranzo, J., Koonin, E. V, Prangishvili, D. & Krupovic, M. Bipartite network analysis of the archaeal virosphere: evolutionary connections between viruses and capsid-less mobile elements. J. Virol. 90, 11043–11055 (2016).

26. Prangishvili, D., Bamford, D. H., Forterre, P. & Iranzo, J. The enigmatic archaeal virosphere. Nat. Rev. Microbiol. 15, 724–739 (2017).

27. Krupovic, M., Cvirkaite-Krupovic, V., Iranzo, J., Prangishvili, D. & Koonin, E. V. Viruses of archaea: Structural, functional, environmental and evolutionary genomics. Virus Res. 244, 181–193 (2018).

28. Garushyants, S. K., Kazanov, M. D. & Gelfand, M. S. Horizontal gene transfer and genome evolution in Methanosarcina. BMC Evol. Biol. 15, 1–14 (2015).

29. Mavrich, T. N. & Hatfull, G. F. Bacteriophage evolution differs by host, lifestyle and genome. Nat. Microbiol. 2, 17112 (2017).

30. Krupovic, M., Prangishvili, D., Hendrix, R. W. & Bamford, D. H. Genomics of bacterial and archaeal viruses: dynamics within the prokaryotic virosphere. Microbiol. Mol. Biol. Rev. 75, 610–35 (2011).

31. Iranzo, J., Krupovic, M. & Koonin, E. V. The double-stranded DNA virosphere as a modular hierarchical network of gene sharing. MBio 7, e00978–16 (2016).

32. Rosvall, M. & Bergstrom, C. T. Multilevel compression of random walks on networks reveals hierarchical organization in large integrated systems. PLoS One 6, (2011).

33. Wolf, Y. I. et al. Origins and Evolution of the Global RNA Virome. mBioBio 9, 1–31 (2018).

34. Koonin, E. V, Dolja, V. V & Krupovic, M. Origins and evolution of viruses of eukaryotes: The ultimate modularity. Virology 479–480, 1–24 (2015).

35. Song, S. & Wood, T. K. Post-segregational killing and phage inhibition are not mediated by cell death through toxin/antitoxin systems. Front. Microbiol. 9, 1–6 (2018).

36. Marraffini, L. A. CRISPR-Cas immunity in prokaryotes. Nature 526, 55–61 (2015).

37. Borges, A. L., Davidson, A. R. & Bondy-Denomy, J. The Discovery, Mechanisms, and Evolutionary Impact of Anti-CRISPRs. Annu. Rev. Virol. 4, 1–23 (2017).

38. Mochimaru, H. et al. Methanolobus profundi sp. nov., a methylotrophic methanogen isolated from deep subsurface sediments in a natural gas field. Int. J. Syst. Evol. Microbiol. 59, 714–718 (2009).

39. O’Leary, N. A. et al. Reference sequence (RefSeq) database at NCBI: Current status, taxonomic expansion, and functional annotation. Nucleic Acids Res. 44, D733–D745 (2016).

40. Brister, J. R., Ako-Adjei, D., Bao, Y. & Blinkova, O. NCBI viral Genomes resource. Nucleic Acids Res. 43, D571–D577 (2015).

41. Delcher, A. L., Bratke, K. A., Powers, E. C. & Salzberg, S. L. Identifying bacterial genes and endosymbiont DNA with Glimmer. Bioinformatics 23, 673–679 (2007).

42. Edgar, R. C. MUSCLE: a multiple sequence alignment method with reduced time and space complexity. BMC Bioinformatics 5, 113 (2004).

43. Remmert, M., Biegert, A., Hauser, A. & Söding, J. HHblits: lightning-fast iterative protein sequence searching by HMM-HMM alignment. Nat. Methods 9, 173–175 (2011).

44. Fu, L., Niu, B., Zhu, Z., Wu, S. & Li, W. CD-HIT: Accelerated for clustering the next-generation sequencing data. Bioinformatics 28, 3150–3152 (2012).

45. Eddy, S. R. Accelerated Profile HMM Searches. PLoS Comput. Biol. 7, e1002195 (2011).

46. Petersen, T. N., Brunak, S., Von Heijne, G. & Nielsen, H. SignalP 4.0: Discriminating signal peptides from transmembrane regions. Nat. Methods 8, 785–786 (2011).

47. Krogh, A., Larsson, B., von Heijne, G. & Sonnhammer, E. L. L. Predicting transmembrane protein topology with a hidden Markov model: application to complete genomes. J. Mol. Biol. 305, 567–80 (2001).

48. Camacho, C. et al. BLAST+: architecture and applications. BMC Bioinformatics 10, 421 (2009).

49. Liaw, A. & Wiener, M. Classification and Regression by randomForest. R News 2, 18–22 (2002).

50. Strobl, C., Boulesteix, A. L., Kneib, T., Augustin, T. & Zeileis, A. Conditional variable importance for random forests. BMC Bioinformatics 9, 1–11 (2008).

51. Simon, N., Friedman, J., Hastie, T. & Tibshirani, R. Regularization paths for Cox’s proportional hazards model via coordinate descent. J. Stat. Softw. 39, 1–13 (2011).

52. Wickham, H.. ggplot2: Elegant Graphics for Data Analysis. (Springer Publishing Company, 2016).

53. R Core Team. R: A Language and Environment for Statistical Computing. (R Foundation for Statistical Computing, 2018).

54. Adriaenssens, E. M., Krupovic, M. & Knezevic, P. Taxonomy of prokaryotic viruses: 2016 update from the ICTV bacterial and archaeal viruses subcommittee. Arch. Virol. 162, 1153–1157 (2017).

55. Kurtz, S. et al. Versatile and open software for comparing large genomes. Genome Biol. 5, R12 (2004).

56. Alva, V., Nam, S.-Z., Söding, J. & Lupas, A. N. The MPI bioinformatics Toolkit as an integrative platform for advanced protein sequence and structure analysis. Nucleic Acids Res. 44, W410–W415 (2016).

57. Demchak, B. et al. Cytoscape: the network visualization tool for GenomeSpace workflows. F1000Research (2014). doi:10.12688/f1000research.4492.2

58. Muhire, B. M., Varsani, A. & Martin, D. P. SDT: A virus classification tool based on pairwise sequence alignment and identity calculation. PLoS One 9, (2014).

59. Eloe-Fadrosh, E. A. et al. Global metagenomic survey reveals a new bacterial candidate phylum in geothermal springs. Nat. Commun. 7, 1–10 (2016).

60. Yu, F. B. et al. Microfluidic-based mini-metagenomics enables discovery of novel microbial lineages from complex environmental samples. Elife 6, 1–20 (2017).

61. Parks, D. H. et al. Recovery of nearly 8,000 metagenome-assembled genomes substantially expands the tree of life. Nat. Microbiol. 2, 1 (2017).

62. Tatusov, R. L. The COG database: a tool for genome-scale analysis of protein functions and evolution. Nucleic Acids Res. 28, 33–36 (2000).

63. Katoh, K. & Standley, D. M. MAFFT multiple sequence alignment software version 7: Improvements in performance and usability. Mol. Biol. Evol. 30, 772–780 (2013).

64. Criscuolo, A. & Gribaldo, S. BMGE (Block Mapping and Gathering with Entropy): A new software for selection of phylogenetic informative regions from multiple sequence alignments. BMC Evol. Biol. 10, (2010).

65. Price, M. N., Dehal, P. S. & Arkin, A. P. FastTree 2--approximately maximum-likelihood trees for large alignments. PLoS One 5, e9490 (2010).

66. Mukherjee, S. et al. Genomes OnLine Database (GOLD) v.6: Data updates and feature enhancements. Nucleic Acids Res. 45, D446–D456 (2017).

67. Hunter, J. D. MATPLOTLIB: A 2D GRAPHICS ENVIRONMENT. Comput. Sci. Eng. 9, 90–95 (2007).

68. Capella-Gutiérrez, S., Silla-Martínez, J. M. & Gabaldón, T. trimAl: A tool for automated alignment trimming in large-scale phylogenetic analyses. Bioinformatics 25, 1972–1973 (2009).

69. Nguyen, L. T., Schmidt, H. A., Von Haeseler, A. & Minh, B. Q. IQ-TREE: A fast and effective stochastic algorithm for estimating maximum-likelihood phylogenies. Mol. Biol. Evol. 32, 268–274 (2015).

70. Letunic, I. & Bork, P. Interactive tree of life (iTOL) v3: an online tool for the display and annotation of phylogenetic and other trees. Nucleic Acids Res. 44, W242–5 (2016).

71. Krupovic, M. Networks of evolutionary interactions underlying the polyphyletic origin of ssDNA viruses. Curr. Opin. Virol. (2013). doi:10.1016/j.coviro.2013.06.010

72. Carr, S. B., Phillips, S. E. V. & Thomas, C. D. Structures of replication initiation proteins from staphylococcal antibiotic resistance plasmids reveal protein asymmetry and flexibility are necessary for replication. Nucleic Acids Res. 44, 2417–2428 (2016).

73. Finn, R. D. et al. The Pfam protein families database: Towards a more sustainable future. Nucleic Acids Res. 44, D279–D285 (2016).

74. Kelley, L. A., Mezulis, S., Yates, C., Wass, M. & Sternberg, M. The Phyre2 web portal for protein modelling, prediction, and analysis. Nat. Protoc. 10, 845–858 (2015).

75. Letunic, I. SMART 4.0: towards genomic data integration. Nucleic Acids Res. 32, 142D–144 (2004).

76. Makarova, K. S. et al. An updated evolutionary classification of CRISPR-Cas systems. Nat. Rev. Microbiol. 13, 722–736 (2015).

77. Hill, J. T. et al. Poly peak parser: Method and software for identification of unknown indels using sanger sequencing of polymerase chain reaction products. Dev. Dyn. (2014). doi:10.1002/dvdy.24183.

78. Lanfear, R. sangeranalyseR: a suite of functions for the analysis of Sanger sequence data in R. (2015).

79. Wickham, H., Hester, J. & Francois, R. readr: Read Rectangular Text Data. (2017).

80. Waterhouse, A. M., Procter, J. B., Martin, D. M. A., Clamp, M. & Barton, G. J. Jalview Version 2--a multiple sequence alignment editor and analysis workbench. Bioinformatics 25, 1189–1191 (2009).

81. Oberortner, E., Cheng, J. F., Hillson, N. J. & Deutsch, S. Streamlining the Design-to-Build Transition with Build-Optimization Software Tools. ACS Synth. Biol. 6, 485–496 (2017).

82. Qiu, D., Damron, F. H., Mima, T., Schweizer, H. P. & Yu, H. D. PBAD-based shuttle vectors for functional analysis of toxic and highly regulated genes in Pseudomonas and Burkholderia spp. and other bacteria. Appl. Environ. Microbiol. 74, 7422–7426 (2008).

