## Supplementary Text and Figures for "Cryptic inoviruses are pervasive in bacteria and archaea across Earth’s biomes"

Identification of putative marker genes and characteristic features for inovirus detection

Genome sequences and predicted proteins from 56 reference *Inoviridae* genomes (Table S1) were gathered, and their predicted proteins were grouped into protein families using (i) all-vs-all blast and InfoMap[1] to define protein clusters, and (ii) HHSearch[2] to combine these clusters into larger protein families (see Methods). A bipartite network was then built using genomes and protein families as nodes, and connecting genomes to protein families when at least one protein affiliated to this family was encoded in the genome. The topology of this network was found to correctly recapitulate the known *Inoviridae* taxonomy, as well as their known host range (Figure S1A). This network was thus used to identify putative core genes that could be used as marker to search for inovirus sequences across new datasets. These core genes would appear in the network as protein family nodes connecting to a maximum of genomes (Figure S1A).

No protein family was universally detected in all the reference genomes, and only morphogenesis proteins (pI) were good candidates for a marker gene: these proteins were split into 3 families only, except for the pI protein from Acholeplasma virus MV-L1 which was a singleton. Two of these protein families included sequences currently annotated as pI and displayed significant hits to the Zot PFAM domain (the only PFAM domain including pI-like proteins), while the third was identified as a pI-like protein based on its unique presence in all *Vespertilliovirus* genomes, its size being consistent with known pI proteins, and a low similarity to the Zot PFAM domain (hhsearch score of 15 and e-value of 0.004), while no other proteins in these genomes displayed any similarity to Zot. Eventually, the complete set of marker included these 3 protein families, the PFAM Zot domain, as well as the putative pI from Acholeplasma phage MV-L1. For each new family, HMM profiles were generated as follows: sequences were first clustered at 90% AAI with cd-hit[3], then aligned with muscle[4], and the hmm profile built with hmmbuild[5].

Canonical *Inoviridae* major coat proteins could be detected based on their length (30 to 90 aa, Table S1), and the presence of a single transmembrane domain (TMD). A signal peptide was detected in most of these proteins (21 of 30) and the corresponding sequences had to be matured in silico (i.e. the signal peptide removed) to recover the expected size and single TMD. About half of minor coat proteins (29 of 49) could also be identified using the same features, but the remaining minor coat were either shorter (e.g. NP_039618) or longer (e.g. YP_002925193). Notably, a good major coat candidate, i.e. a protein of 30 to 90 aa and with 1 TMD, was detected in every *Inoviridae* genome, even the ones for which no protein was annotated as a major coat (Table S1).

Design of an automatic classifier to detect inovirus sequences

To automatically detect inovirus genomes, we first searched for new inovirus sequences to add to the 56 genomes database, in order to gather a positive dataset large enough for training an automatic classifier. To that end, we used the set of pI HMM profiles previously described (see above) to search 56,868 bacterial and archaeal genomes publicly available in the IMG database, which yielded 6,819 hits (hmmsearch[5], score ≥ 30 E-value ≤ 0.001). The genomic context of these pI-like genes was then examined in a window of 20 genes in 5' and 3' by (i) gathering the PFAM annotation of the genes in these regions from IMG, (ii) affiliating these genes to the previously generated reference *Inoviridae* protein families (hmmsearch[5], thresholds of 30 on score and 0.001 on e value), and (iii) predicting putative *Inoviridae* coat proteins based on protein size, presence of a signal peptide, and single TMD (see above). From these annotations, putative complete inovirus genomes were identified by extending the prediction around the initial pI protein in 5' and 3' until reaching a protein affiliated to a PFAM domain never encountered in a reference *Inoviridae* genome (i.e. "unexpected PFAM affiliation"), and then assessing if the corresponding prediction either (i) spanned an entire circular contig with an expected inovirus genome size (i.e. 5-20kb), or (ii) included putative canonical attachment (att) site, i.e. direct repeats of 10bp or longer that could be identified in a tRNA gene or directly outside of an integrase gene[6]. A total of 795 putative inovirus genomes were detected: 213 as circular contigs, and 582 as integrated prophages. Their predicted pI proteins were added to the references to generate new improved HMM models, and another round of search of the same datasets was conducted, adding an additional 10 putative genomes (3 circular, 7 prophages with canonical att site). The gene content of these genomes was next manually inspected to verify that these were consistent with known *Inoviridae*, and edge cases were excluded.

Next, the *Inoviridae* reference isolates and these 805 manually curated sequences were gathered as a positive set to train an automatic classifier, in order to be able to automatically evaluate a putative inovirus genome detected based on the presence of a pI-like gene. A negative set was generated by taking random fragments in genomes where an inovirus sequence was detected (n=1,000), as well as genome fragments around pI proteins manually identified as false positives, i.e. not inovirus sequences (n=1,000), with a fragment length following the length distribution of the complete inovirus genomes in both cases. This was done to ensure that the model is trained on negative cases representing both typical genome fragments from inovirus hosts, as well as typical genome context for pI-like proteins which are not associated with inovirus prophages.

A set of genome features was identified that could be used to identify genuine inoviruses (see examples in Figure S1B). These include (i) fragment length, (ii) number of genes in the fragment, (iii) number of proteins with a hit to a pI protein family, (iv) number of genes with a significant hit to inovirus capsid PFAM domains, (v) number of genes predicted as inovirus coat proteins, (vi) number of genes with a significant hit to reference *Inoviridae* protein families, (vii) number of genes without an unexpected PFAM affiliation, (viii) percentage of genes in the fragment without an unexpected PFAM affiliation, (ix) median gene length in the fragment, and (x) first decile of gene length in the fragment. "Expected" affiliations were based on the affiliation of known inovirus proteins to PFAM domains and to their associated keywords ("DUF", "HTH", "DNA", "repeat", "toxin", and "regul"), while "Unexpected" PFAM domains are the rest of the PFAM database.

Based on the training sets, a random forest classifier was found to be the most efficient at discriminating inoviruses from background host genome (compared to random forest with conditional inference and generalized linear model with lasso regularization), and achieved (at the selected threshold of score $\geq 0.9$) 92.5% recall (percentage of "true" inoviruses correctly predicted as inoviruses), 99.9% specificity (percentage of "true" non-inoviruses sequences correctly predicted as non-inoviruses), and 99.8% precision (percentage of "true" inoviruses within sequences predicted as inoviruses). This model and threshold combination was chosen because it provided the maximum recall at the low false discovery rate of 0.2% (Figure S1C). This approach can thus be used in place of the manual curation step to evaluate genome regions surrounding putative pI proteins, and systematically detect new inovirus sequences with high accuracy.

Identification of non-inovirus ATPases among putative pI proteins

When detecting new inoviruses using pI-like proteins, the presence of an ATPase domain in these proteins can lead to false positive detections. Our automatic classifier is able to identify most of these non-inovirus ATPases based on genome context, as illustrated by the large number of hits for which no inovirus genome was predicted (gray sections of the pie charts in Fig. 1D). However some false positives may remain, for instance due to another virus or mobile genetic element with atypical genes and/or short geness encoding a related ATPase. To identify and remove these sequences, we explored the protein clusters (PCs) computed from the complete set of genes from all inovirus species (both known and newly detected), and examined the ones including at least one protein initially identified as pI at the first detection step, i.e. used as a starting point for the detection of inovirus sequence. Overall, 6,570 non-redundant proteins were detected across 45 different pI-like protein clusters (PCs), with 16 singletons. These 45 different PCs gathered into 10 different iPFs ("inovirus Protein Families", Figure S2A, see Methods). Two of these iPFs (iPF_00003 and iPF_00013) included a large number of mostly (> 95%) pI-like proteins (4,548 and 1,740 proteins, respectively). The few sequences in these iPFs that had not been previously identified as pI-like protein were usually genuine partial pI-like proteins too short to yield a significant hit, and thus unaffiliated. Both of these iPFs also included pI proteins from *Inoviridae* isolates, and were thus annotated as genuine pI proteins. The multiple alignments of the 34 PCs clustered into these 2 iPFs were visually inspected to verify that (i) the ATPase domain was most closely related to the inovirus Zot domain as opposed to one of the other known FtsK/HerA ATPase domains in the PFAM database (PF01580, PF01935, PF02534, PF03135, PF05872, PF06834, PF09378, PF09397, PF10412, PF11130, PF12538, PF12696, PF12846, and PF13491), and (ii) the sequence included a TMD to anchor the pI protein in the host membrane.

The 21 PCs clustered in iPF_00003 corresponded to sequences with a "typical" pI protein architecture, i.e. the ATPase domain is followed by a C-terminal extension with a TMD membrane (Figure S2B). These PCs were mostly associated with gram negative hosts (with the exception of some prophages detected in Clostridia), and included all gram negative *Inoviridae* isolates (e.g. M13, CTX, etc). However, 3 PCs were identified that could not be confirmed as likely inovirus pI protein: PC_00610, PC_01272, and PC_01338 were all most similar to the ATPase domain from archaeal turrivirus STIV, and were thus considered as false positives and removed from the inovirus dataset.

Conversely, the 13 PCs clustered in iPF_00013 displayed an "atypical" pI protein architecture: their Zot domain was not followed by a C-terminal extension, and the TMD was usually detected in the N-terminal part of the protein (or in the case of PC_00303 at the C-terminal tip of the sequence). These PCs are mostly composed of sequences found in gram positive hosts, and include *Inoviridae* isolated on *Propionibacterium*, *Thermus*, and *Spiroplasma* (Figure S2A, Table S3). The distribution of pI-like proteins in two distinct iPFs thus seems to be roughly correlated with the fundamental differences between cell membranes of gram negative hosts, associated with typical pI, and cell membranes of gram positive or wall-less hosts, associated with atypical pI. Among the PCs gathered into iPF_00013, one (PC_01836) was identified as likely false positive as it was most closely related to archaeal turrivirus STIV, and the associated sequences were excluded from the inovirus dataset.

Finally, for pI-like proteins outside of iPF_00003 and iPF_00013, genomes were individually inspected to evaluate whether these could also represent inovirus genomes. All but 1 of these sequences were identified as likely false positives based on the similarity of their pI-like sequence to another FtsK/HerA ATPase domain with higher scores than to the Zot domain. The only case of a putative genuine inovirus pI protein found outside of iPF_00003 and iPF_00013 was sequence 1066081_contig_758_11 found in iPF_00002. This iPF was annotated as an assembly protein, and the clustering of this specific sequence seemed to originate from a fusion of the pI (Morphogenesis) and pIV (Assembly) proteins (Figure S2B). Interestingly, this potential fusion of pI and pIV genes has also been identified in PC_01246, annotated as a pI-like protein (part of iPF_00003). Notably, no other pI or pIV were identified in the genomes encoding these putative fusion proteins, and these sequences clearly included both conserved domains (Figure S2B). In characterized inoviruses, the assembly domain (pIV) is encoded by a distinct gene and ensures the passage of the virion across the outer membrane in diderm hosts. Hence, these fused genes could produce a protein which would allow the passage of the virion across both host membranes, and accordingly these were all detected in diderm hosts. Although very atypical, these sequences including fused pI-pIV proteins were still included in the final dataset as they are likely functioning extrusion mechanisms, at least based on sequence analysis. Conversely, 5 other putative pI-pIV fusion proteins were detected in iPF_00002, but displayed a seemingly truncated Zot-like domain and lacked the typical TMD found in C-terminal of this Zot-like domain. These sequences were not considered as genuine pI-like proteins.

Eventually, this improved annotation of pI proteins was used to refine the final dataset of inovirus sequences. This included 4 sequences initially identified as "tandem prophages" which were reclassified as "regular genomes" since one of the two pI detections was a false positive, and 28 sequences removed from the dataset because their putative pI protein was found in one of the false positive PCs.

Types of inovirus sequences detected

Among the new inovirus sequences detected, 1,709 were identified as putative complete inovirus genomes as these were either circular contigs (n=1,088), prophages with identified canonical attachment (att) site in a tRNA (n=311) or prophages with an identified canonical att site adjacent to an integrase-like gene (n=310, see Methods). An additional 1,586 fragments were putative complete prophages for which non-canonical att sites could be identified, i.e. the fragment is framed by direct repeats but these repeats are not within a tRNA gene or outside an integrase-like gene. Finally, the remaining fragments were either linear contigs likely from partial genomes (n=2,526) or prophages for which no att site could be identified (n=4,474). Notably, 553 fragments included multiple distinct pI-like proteins with no identifiable genome ends or attachment sites, and as such likely represent tandem prophage insertions, including possibly degraded prophages[7].

Distribution of inovirus sequences across metagenomes and biomes

The 5,917 inovirus sequences detected in metagenome assemblies, which included 3,677 species exclusively detected from metagenomes (Figure S1D, Table S3), can inform about the distribution of these viruses across ecosystems and geographic locations. Overall, individual species tend to be associated with a single sample type: 95% of species detected in multiple metagenomes are restricted to a single sample type (Table S3). Inovirus sequences were detected in environments ranging from mesophilic (e.g. freswhater lakes) to 'extreme' (e.g. thermal springs or deep-ocean subsurface), from pristine (e.g. Antarctica) to strongly impacted by human activity (e.g. wastewater), from free-living microbial communities (e.g. ocean surface) to host-associated (e.g. human gut, rhizosphere), as well as on every continent and from the equator to the poles. Associated with their broad host range, the extensive ecological distribution of inovirus sequences suggests they have the potential to impact most of Earth's ecosystem, including modulating interactions between organisms within holobionts, as suggested by some available isolates[8].

Prevalence of inoviruses in microbial genomes

Based on the 2,289 inovirus species associated to a host, we calculated an estimated prevalence for inoviruses, i.e. the proportion of microbial genomes including an inovirus genome. The highest median prevalence was observed in Gamma- and Betaproteobacteria, where qualified genera (i.e. genera with $\geq$ 5 genomes) displayed on average > 10% of genomes with $\geq$ 1 detection(s) (Figure S3A). Among these, prevalence in the genus *Xylella* was particularly high (87%), although these prophages were all associated with the microbial pathogen *Xylella fastidiosa*, and thus likely reflect the strong association of inoviruses with this specific species. Beyond these two groups, inoviruses were detected in ~1% of genomes on average, although this prevalence was > 15% for 5 genera (*Acidithiobacillus*, *Desulfosporosinus*, *Eubacterium*, *Lachnoclotridium,* and *Spiroplasma*), suggesting these might be evolving under an unusually high inovirus infection rate (Figure S3A).

Curiously, 5 host genera composed of > 400 genomes did not yield any detection: *Mycobacterium* and *Streptomyces* (Actinobacteria, n=660 and 411, respectively), *Helicobacter* (Campylobacterota, n=443), *Lactobacillus* and *Staphylococcus* (Firmicutes, n=692 and 844 respectively). Since filamentous phages have been detected in other members of the same families, it is likely that members of these specific genera are very rarely (if ever) infected by inoviruses.

Co-infection patterns across host groups

Usually, a single inovirus sequence was detected per genome (76% of cases), and multiple detections were mostly found within the two host classes with high inovirus prevalence (Gamma- and Beta-proteobacteria, Figure S3B). However, *Spiroplasma* represented an exception to this rule: beyond a unusually high level of inovirus prophages compared to other Tenericutes, these genomes also displayed an average of > 15 distinct detections, including 2 genomes with 24 and 25 distinct prophages detected (Table S3). Although these data are only based on 5 *Spiroplasma* genomes in which prophages were detected, it suggests that at least some members of this genus may be uniquely able to integrate and maintain dozens of distinct inovirus genomes at a time, a feature previously hypothesized as driving the extensive intra-genome recombinations observed in this clade[9].

Inovirus prophages were frequently detected along with *Caudovirales* prophages: 1,573 bacterial genomes included signs of both types of viruses, consistent with a trend previously noted in smaller scale prophage analyses[10,11] (Figure S3C). Curiously, these combined prophages insertions sometimes occurred at the same location in the host genome, particularly in Betaproteobacteria and Campylobacterota such as *Neisseria* and *Campylobacter* (Figure S3D & E). Such co-localization could provide opportunity for horizontal gene transfer through imprecise excision, and more generally highlights a potential for direct virus-virus interactions, which have so far remained mostly unexplored[11].

Evaluation of the taxonomic rank represented by network-derived genome (sub)groups

ICTV guidelines are only available for genera (75% AAI) and species (95% ANI) in the *Inoviridae* family, such that we had to use other viral groups as reference to estimate which taxonomic rank the new groups and sub-groups defined based on gene content comparison represented. To this end, we compared the Amino Acid Identity percentage (AAI) of marker genes (i.e. pI-like proteins) from this new sets of inovirus genomes with other established viral groups at different ranks. For the order rank, we opted to use *Caudovirales* as references even though these are dsDNA viruses and tend to have larger genomes, since no classification at the order rank is available for small ssDNA viruses. For family and genus, we used established ssDNA taxonomy from the *Microviridae* and *Circoviridae* families, more comparable to inoviruses in terms of genome size and complexity.

This comparison to known viral taxonomic groups suggested that the 6 main groups observed on the inovirus sequence network are comparable to currently established viral families, while levels of similarity observed when comparing sequences between the 6 main groups were consistent with an order rank (Figure S5B). Hence we propose that the *Inoviridae* family should be considered as a viral order instead, which we would propose to name *Inovirales* in accordance with standards in viral taxonomy nomenclature. This order would be tentatively divided into 6 candidate families, corresponding to the 6 mains groups established from the genome-PC network. Still based on AAI, the 212 sub-groups would be consistent with subfamilies, as these are more divergent than the established threshold used to define genera in the current *Inoviridae* family (75% AAI[12]) and more divergent than currently established ssDNA virus genera (Figure S5B). As would be expected, all members of each current genus were found in a single proposed subfamily (Table S3).

<u>Genome network topology and connector PCs</u>

Overall, only 20 PCs (out of 892 displayed on the network) connected genomes across proposed families (Figure 5). All but 3 of these (i.e. 85%) were functionally affiliated, including 4 pI-like, 3 structural, and 7 replication-associated proteins, suggesting these "connector" PCs (*sensu* [13]) likely represent some of the most conserved genes across inovirus genomes. None of these PCs however connected substantially (>50% proposed subfamilies) to more than 1 proposed family, suggesting that these most likely reflect events of horizontal gene transfer or convergent evolution involving these conserved genes. This further illustrates the complex evolutionary history of these genomes for which no single gene seems to be both conserved and exclusively (or near-exclusively) vertically inherited (Figure S5A).

<u>Characteristics and proposed names for new proposed families</u>

The two largest proposed families (in dark blue and teal in Figure 5) comprise 3,576 and 1,020 genomes, respectively, and include all isolates officially classified into the seven *Inoviridae* genera currently recognized by the ICTV (Table S3). The first of these two proposed families (in teal in Figure 5) includes members of genera *Fibrovirus, Habenivirus, Inovirus, Lineavirus* and *Saetivirus*, and so gather the prototypical and most characterized isolated *Inoviridae.* The second one (dark blue) comprises members of the genus *Vespertilinovirus* and the single member of the *Plectrovirus* genus. Hereinafter, we refer to these two putative families as "Protoinoviridae" (for the inovirus prototypical members) and "Vespertilinoviridae" (inspired by the main genus of this proposed family), respectively. The proposed "Protoinoviridae" family comprises genomes nearly exclusively associated with Gamma- and Beta-proteobacteria, while the "Vespertilinoviridae" include mostly genomes associated with Clostridia and Tenericutes (Figure 5, Figure S5D). A third putative family includes the remaining isolates unassigned to a genus yet. These genomes tend to be smaller than those of other inoviruses (median size of 6.1kb), thus, we propose to name this candidate family "Paulinoviridae" (from 'paulus', latin for little/small). "Paulinoviridae" genomes are primarily detected in hosts affiliated to Actinobacteria, CPR, and Deinococcus-Thermus clades (Figure S5D).

The remaining three proposed families do not include any viral isolate, but two of these exhibit specific genome features (Figure S5C & D). The first putative family is composed of large genomes (median 9.4kb); hence we proposed to name this group "Amplinoviridae" (from "amplus", latin for large). The second one is composed of genomes with high coding density (i.e. more genes for comparable genome size, median number of genes=16) and we propose to name this assemblage "Densinoviridae" (Figure S5C). Members of the "Amplinoviridae" are largely associated with hosts from Deltaproteobacteria and Campylobacterota, whereas "Densinoviridae" are predominantly found in Bacilli and Chloroflexi (Figure S5D). Notably, two sequences in the "Densinoviridae" have been previously described as "cryptic plasmids" in Bacilli[14]. Similarities between small plasmids and filamentous phages have long been noted, and the boundary between the two types of mobile genetic elements seems tenuous at best[15]. However, filamentous particles have been induced from similar bacteria[16], and we have identified putative capsid proteins, a hallmark of viruses, encoded by members of the "Densinoviridae" (see main results and Figure S6A). Hence, given that inoviruses have been frequently confused with plasmids (e.g. NC_002473 and NC_010429), these sequences are likely to correspond to genuine novel inovirus genomes. The last proposed family includes the only inoviruses associated with photosynthetic Cyanobacteria, hence we propose to name this candidate family "Photinoviridae".

Although the proposed families were defined exclusively from gene content analysis, they exhibited specific genome features and host ranges which suggested they indeed represent coherent groups. First, proposed families differ in terms of genome size and number of genes predicted. Notably, the median genome size within candidate families varied from 6kb ("Paulinoviridae") to 9.5kb ("Amplinoviridae"), and although most groups encoded a median of 11 to 13 genes per genome, one ("Densinoviridae") displayed a median number of genes of 16 (Figure S5C). In addition, each of the proposed family is associated with a specific host range, with very little overlap (Figure S5D): of the 70 host families with at least 2 inovirus sequences detected, 61 were associated with a single proposed inovirus family, 6 are associated with 2 proposed inovirus families, and only 3 are associated with 3 proposed inovirus families (*Peptococcaceae*, *Paenibacillaceae*, and *Bacillaceae*, all in the Firmicutes phylum, Table S3).

Contrasting with their host range, the proposed inovirus families were not structured by biome or ecosystem type (Figure S5D). All six candidate families seem to be detected in virtually every type of environment, and the cases of non-detection are associated with under-sampled groups (i.e. proposed families with < 400 species). These data can however point toward which specific biome to sample in priority when targeting individual candidate families: "Vespertilinoviridae" and "Amplinoviridae" seem to be enriched in human-associated samples, "Densinoviridae" in "extreme" aquatic environments such as deep sub-surface, thermal spring, and hypersaline lakes, while both "Paulinoviridae" and "Photinoviridae" are preferentially detected in soil samples.

<u>Identification and annotation of putative archaea-associated inoviruses</u>

The four putative proviruses were identified in the genome sequences of three isolates, two affiliated to *Methanolobus* and one to *Methanosarcina*, all in the *Methanosarcinacea* family of the phylum Euryarchaeota, as well as one metagenome-assembled genome (MAG) affiliated to the Aenigmarchaeota candidate phylum. The contigs composing this MAG were inspected to confirm that they represented a single and cohesive population genome, and no sign of contamination, i.e. presence of a contig affiliated to a different microbial genome, could be identified. The gene content of these different inoviruses was consistent with their respective host: the 3 *Methanosarniacea*-associated viruses displayed little to no sequence similarity to the sequence detected in the Aenigmarchaeota MAG (Figure 4A).

The two sequences detected in *Methanolobus* included the full repertoire of genes expected in a genuine inovirus, including a morphogenesis (pI) protein with an N-terminal TMD typical of inoviruses infecting monoderm hosts, an integrase gene, genes predicted to encode structural proteins based on sequence length and presence of a single TMD, as well as a gene encoding a rolling-circle replication initiation protein[17]. This gene complement strongly suggests that these two sequences represent fully functional inoviruses, since they include the full suite of genes required for genome integration, replication, encapsidation, and extrusion (Figure 4A). The detection of genes predicted as structural, i.e. short genes with a single TMD, is especially noticeable given that only 0.69% of all genes in Euryarchaeota display features characteristic of structural proteins of inoviruses (30-90 aa, 1 TMD). The predicted proviruses are thus more likely to be inoviruses than any other type of mobile genetic element.

The putative proviruses identified in *Methanosarcina* displayed genes for integration, morphogenesis, and predicted structural proteins, but no recognizable gene involved in genome replication. Similarly, the sequence identified in the Aenigmarchaeota MAG only included a morphogenesis gene and a predicted structural protein. Hence, these two latter sequences could be partial genomes, possibly remnants from a decaying provirus, or could be complete genomes of active viruses for which replication-associated gene(s) cannot yet be identified, as is common amoung archaeal viruses[18]. Regardless of the completeness of these genomes, both include an inovirus-like morphogenesis protein suggesting these are most likely inoviruses. In addition, we found a perfect match between the Aenigmarchaeota provirus and a CRISPR spacer from a different contig in the same MAG, which is also consistent with it being a provirus (Table S6).

PCR validation of excision for an inovirus integrated in *Methanolobus profundi* MobM

Attempts at observing inovirus capsids through TEM were unsuccessful because *Methanolobus* MobM flagella are similar in structure, length, and width to filamentous virions[19]. Thus, we used instead PCR to detect the presence of a circularized form of the provirus, which would correspond to the complete genome being excised and replicated or encapsidated (Figure 4B). We first verified that the genome sequencing and assembly was correct by amplifying a product internal to the predicted provirus and a product spanning the predicted insertion site (Figure 4B). In both cases, we obtained a successful amplification with products of the expected size, confirming that the predicted provirus is present and likely integrated in most cells in the culture. Notably, we obtained positive amplification for the product spanning the insertion site from the fraction < 0.22 µm, which suggests that some MobM cells can pass through the 0.22 µm filter typically used to separate viruses from their bacterial or archaeal hosts.

Next, we designed a PCR primer pair specific to the predicted excised form of the virus genome by combining a forward primer from the 3' end of the provirus to a reverse primer in the 5' end of the provirus (Figure 4B). We obtained a product of the expected size, and sequencing of the product confirmed that it spanned both ends of the predicted provirus in the predicted orientation and at the expected coordinates. This latter PCR reaction initially generated more nonspecific products than the internal or integration site primers, and the reaction annealing temperature had to be increased (> 56°C) to obtain a single band at the expected size. This higher level of nonspecific amplification combined with the fact that the product obtained yielded a relatively faint band (Figure 4B) suggests that the template for this reaction, i.e. the excised form of the virus genome, is found in a much smaller fraction of cells than the integrated form. It is thus very likely that under laboratory conditions, even after treatment with mitomycin C, the provirus is repressed in most cells resulting in an overall low concentration of circular virus genomes.

### Additional host associations from CRISPR spacer matches to metagenome-assembled inoviruses

Matches between CRISPR spacers and inovirus sequences included both predicted prophages/ proviruses for which host information could be confirmed (n=711) and metagenome assemblies for which new host information could be obtained (n=439, Table S6). Near-exact matches (i.e. 0 or 1 mismatch) between CRISPR spacers and metagenome-derived viral contigs have been shown to reliably associate uncultivated viral genomes to putative host(s)[20]. Here, the reliability of near-exact CRISPR matches (i.e. allowing at most 1 mismatch over the entire spacer length) was confirmed by the CRISPR-based host links assessed for prophage predictions: in 99.5% of the cases, host affiliations were consistent (708 of 711). The three outliers might be resulting from false positive spacer matches, horizontal virus transfer, or a very broad host range for certain inoviruses. It is of note that introduction of the genome of an inovirus infecting *Clostridium*, a gram-positive bacterium, into the gram-negative *Escherichia coli* resulted in production of filamentous virus-like particles[21], suggesting that host switches might not be strictly prohibited among inoviruses. Nevertheless, the overall agreement between spacer matches and host affiliation of prophages suggest that spacer matches to metagenome-derived inovirus sequences can be used confidently, expanding the number of host-associated sequences to 439 additional putative inovirus species.

Most of the host pairings derived from these metagenome CRISPR spacer matches were found in hosts groups for which prophages had already been detected, and only 4 new orders were identified. First, 2 inovirus sequences were newly associated to *Roseiflexus* genomes in the *Chloroflexales* order. Other sequences from the same phylum (*Chloroflexi*) had been linked to inovirus sequences, and all these *Chloroflexi*-associated sequences were consistently affiliated to the proposed “Densinoviridae” candidate family. In addition, these metagenome-derived inovirus sequences were detected in a hot spring metagenome, consistent with the preferential habitat of *Roseiflexus.*

Another 2 species were newly associated with an *Aphanizomenon* genome (genus of photosynthetic Cyanobacteria). These 2 putative viral sequences were consistently affiliated to the “Photinoviridae” proposed family, which gathers all inoviruses associated with photosynthetic Cyanobacteria, and consistently originated from two freshwater lake metagenomes (sampled from Lake Mendota).

One inovirus species was newly associated with a genome assembled from a *Nasutitermes corniger* (a species of termite) metagenome and currently affiliated as an “Unclassified

Fibrobacteria". No prophage had been detected associated with this specific host phylum so far. Consistently, the inovirus species was also assembled from a termite gut metagenome.

Another inovirus species was newly associated with a genome affiliated to the Nitrospinae phylum-level group. This inovirus species is classified in the "Amplinoviridae" Subfamily 4, which includes Deltaproteobacteria-associated inoviruses. This is consistent with Nitrospinae and Deltaproteobacteria being related groups of bacteria. The inovirus genome was assembled from a groundwater metagenome as was the bacterial genome.

Finally, one species was associated for the first time to *Caldicellulosiruptor obsidiansis*, a host in the *Thermoanaerobacterales* order, part of the *Clostridia* class for which other putative inovirus sequences had been detected. This sequence was consistently affiliated to the proposed subfamily Sf_1 of the "Vespertilinoviridae" candidate family, the main group of Clostridia-infecting inovirus sequences identified in this study, and detected in a hot spring metagenome, which is consistent with the known preferential habitat of *Caldicellulosiruptor obsidiansis*.

<u>Evaluation of hypothetical proteins from self-targeted inoviruses in a *Pseudomonas aeruginosa* model</u>

Hypothetical proteins from two self-targeted *Pseudomonas* inovirus prophages for which no Acr locus could be identified elsewhere in the genomes were synthesized and cloned in a pHERD30T vector for expression in *Pseudomonas aeruginosa* (Figure S7). Two of these candidate genes (2687473922 and 2687473921) were toxic when expressed in the host, and their putative Acr or superinfection exclusion activity could not be assessed. However, these genes may be components of novel toxin-antitoxin systems.

Two candidate genes demonstrated superinfection exclusion activity, which was manifested by the absence of plaques at dilutions for which plaques were formed for the same phage in the same host transformed with the empty vector (Figure S7C). Neither of the 2 genes provided universal superinfection exclusion: gene 2687473927 prevented or limited infection of host strain PAO1 by 3 of the 6 phages tested, but no effect could be observed in the PA14 strain. By contrast, 2687473923 did not provide any superinfection exclusion in host PAO1, but prevented infection of 1 of the 3 phages efficiently infecting PA14 (Figure S7C). This suggests that inovirus-derived superinfection exclusion activity varies depending on the host strain and the co-infecting virus. Specifically, gene 2687473927 seems to have a relatively broad spectrum and could provide a general fitness advantage to host PAO1 by limiting infection in this specific host strain for both temperate Mu-like siphoviruses (DMS3m and JBD30) and lytic T7-like podoviruses (KMV). Conversely, the effect of 2687473923 seems to be much more restricted, and points toward more specific virus-virus interactions or incompatibility between the inovirus and phage JBD30.

Although both proteins are uncharacterized, they are relatively widely distributed in inoviruses, forming two corresponding protein families: iPF_00048 for gene 2687473923 and iPF_00082 for gene 2687473927. Members of the iPF_00048 protein family, responsible for the "narrow" superinfection exclusion, were found in 424 distinct inovirus species. These inoviruses were affiliated across 9 proposed subfamilies within the "Protoinoviridae", and associated with both Beta- and Gammaproteobacteria hosts. Since some members of this protein family contain an HTH domain, we posit that these genes may be coding for transcriptional regulators that could provoke incompatibility with some individual phages, but their primary function might not be superinfection exclusion.

Members of the iPF_00082 protein family ("broad" superinfection exclusion) were detected in 163 distinct inovirus species, all affiliated to the "Protoinoviridae" and nearly all (98%) to the "Protoinoviridae:Sf_2" proposed subfamily. All identified hosts for these species were affiliated to the *Pseudomonas* genus. This narrow distribution in terms of inovirus family/subfamily and host range suggests that members of this protein family have evolved in *Pseudomonas*-specific inoviruses to mediate broad-spectrum superinfection exclusion. Strikingly, nearly half of the inovirus prophages identified in *Pseudomonas* genomes (44%, 158 of 359) encoded this gene. This could be due to positive selection of this gene in inovirus prophages because of its superinfection exclusion properties, although we cannot exclude a potential bias in the *Pseudomonas* genome dataset whereby many strains of *Pseudomonas aeruginosa* with distinct but closely related inovirus prophages would have been sequenced. Finally, all members of the iPF_00082 protein family are 29-30 aa-long and carry predicted α-helical membrane-spanning domain, suggesting that superinfection exclusion may occur at the host cell surface, possibly during the attachment and/or entry of a superinfecting phage. Notably, several *Pseudomonas* dsDNA prophages have already been shown to provide superinfection exclusion through alteration of the host T4 pilus[22], which could be the case as well for these inovirus-encoded proteins.

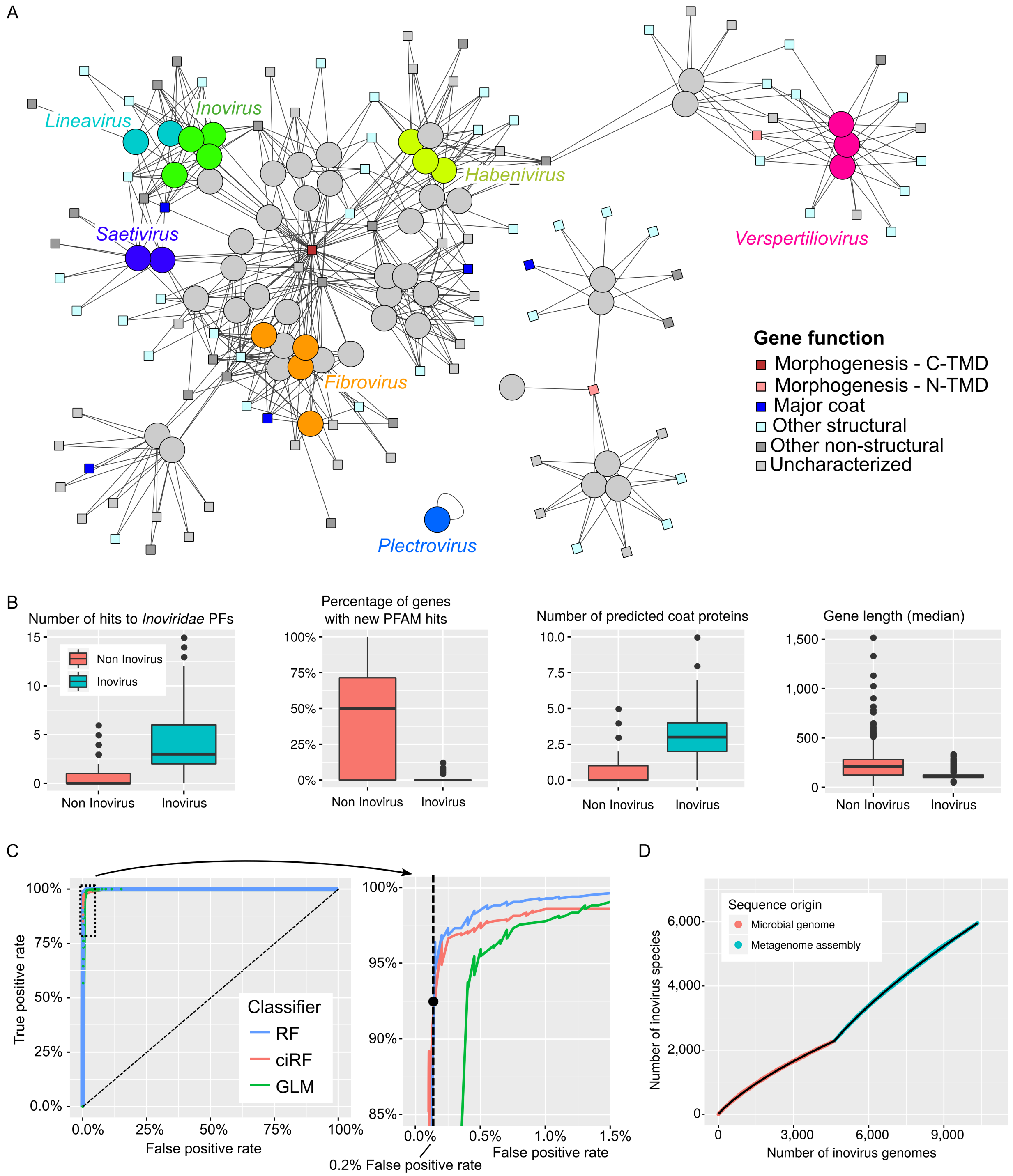


**Figure S1. Identification of marker gene(s) and other characteristic features of inovirus genomes for training of an automatic classifier and recovery of new inovirus species from (meta)genomes.** A. Genome-gene bipartite network of publicly available *Inoviridae* isolate genomes. Genomes are represented as circles colored according to their genus classification, and protein families are displayed as squares colored by their predicted function. ICTV-proposed genera are indicated by coloring of the genome nodes. Morphogenesis (pI-like) proteins are highlighted in red, and although these proteins were represented by 3 distinct protein families, local similarities could still be detected between the corresponding HMM profiles (HHSearch probability ≥ 90%). The two types of pI-like proteins, with the transmembrane domain (TMD) either in C- or N-terminal are indicated in dark and light red respectively (see Figure S2). B. Example of genome features evaluated on isolate inoviruses and manually-curated inovirus prophages (in blue) and other fragments from microbial genomes used in the negative training set (in red). C. ROC curve of the new automatic classifier distinguishing inovirus genomes from other viral or microbial genome fragments. A subplot displays a zoom on the area < 2% false positive rate and > 85% true positive rate. The three types of classifier tested are plotted in different colors, and the true positive and false positive rates associated with the chosen threshold of 0.9 for the random forest classifier indicated with a black circle on the subplot. RF: Random Forest, ciRF: conditional inference Random Forest, GLM: Generalized Linear Model. D. Accumulation curves of inovirus species. The number of different species is indicated as a function of the total number of complete and partial genomes, first for detections in draft and complete genomes from bacteria and archaea, and then in metagenome assemblies. A set of 10 subsample replicates were calculated and are plotted in colors, while the resulting average number of species is plotted in black.

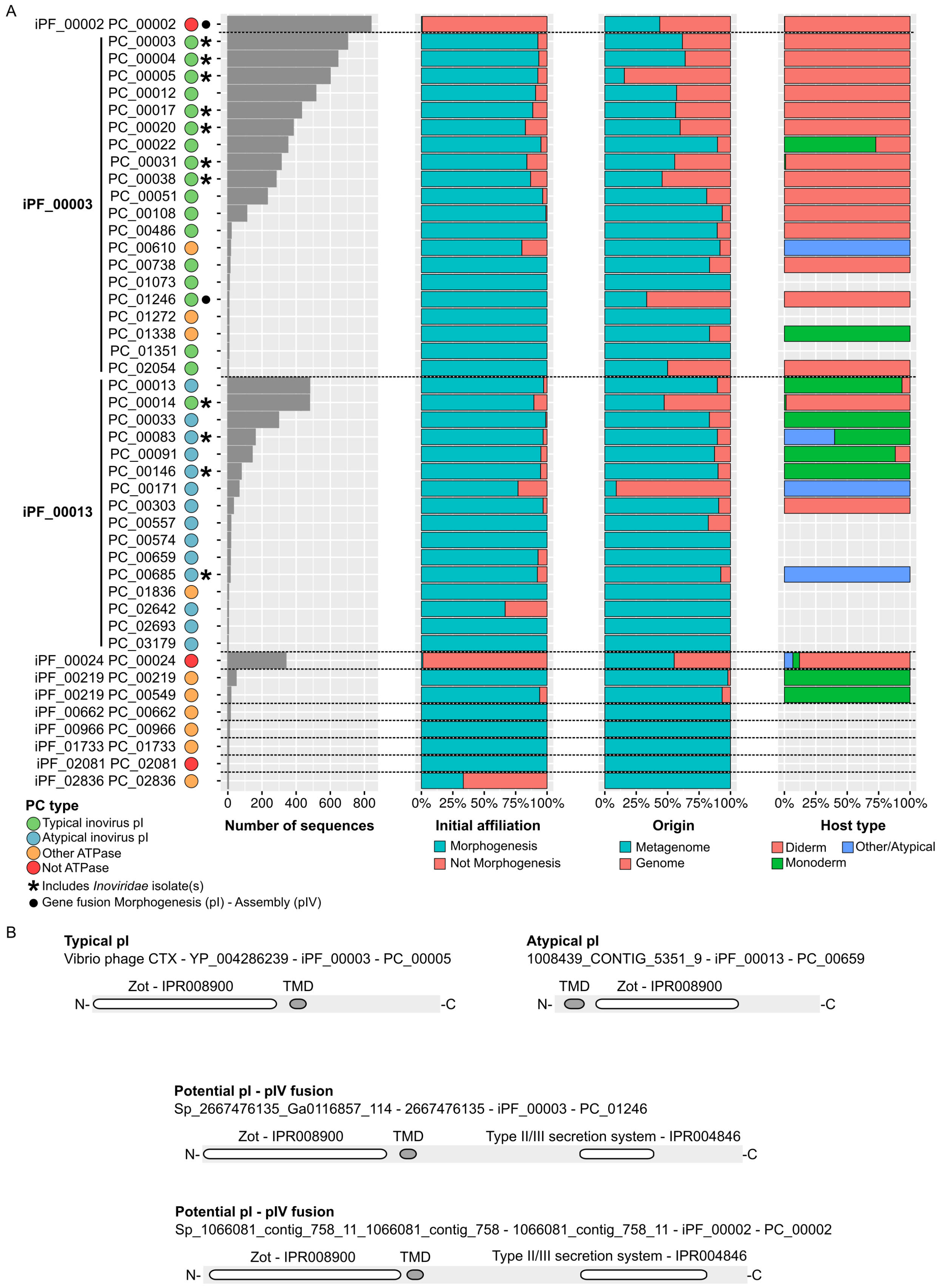
A
iPF_00002 PC_00002
PC_00003
PC_00004
PC_00005
PC_00012
PC_00017
PC_00020
PC_00022
PC_00031
PC_00038
PC_00051
PC_00108
PC_00486
PC_00610
PC_00738
PC_01073
PC_01246
PC_01272
PC_01338
PC_01351
PC_02054
iPF_00003
PC_00013
PC_00014
PC_00033
PC_00083
PC_00091
PC_00146
PC_00171
PC_00303
PC_00557
PC_00574
PC_00659
PC_00685
PC_01836
PC_02642
PC_02693
PC_03179
iPF_00013
iPF_00024 PC_00024
iPF_00219 PC_00219
iPF_00219 PC_00549
iPF_00662 PC_00662
iPF_00966 PC_00966
iPF_01733 PC_01733
iPF_02081 PC_02081
iPF_02836 PC_02836
0 200 400 600 800
Number of sequences
0% 25% 50% 75% 100%
Initial affiliation
Morphogenesis
Not Morphogenesis
0% 25% 50% 75% 100%
Origin
Metagenome
Genome
0% 25% 50% 75% 100%
Host type
Diderm
Other/Atypical
Monoderm
PC type
Typical inovirus pI
Atypical inovirus pI
Other ATPase
Not ATPase
* Includes *Inoviridae* isolate(s)
● Gene fusion Morphogenesis (pI) - Assembly (pIV)
B
Typical pI
Vibrio phage CTX - YP_004286239 - iPF_00003 - PC_00005
Zot - IPR008900
TMD
N-
-C
Atypical pI
1008439_CONTIG_5351_9 - iPF_00013 - PC_00659
TMD
Zot - IPR008900
N-
-C
Potential pI - pIV fusion
Sp_2667476135_Ga0116857_114 - 2667476135 - iPF_00003 - PC_01246
Zot - IPR008900
TMD
Type II/III secretion system - IPR004846
N-
-C
Potential pI - pIV fusion
Sp_1066081_contig_758_11_1066081_contig_758 - 1066081_contig_758_11 - iPF_00002 - PC_00002
Zot - IPR008900
TMD
Type II/III secretion system - IPR004846
N-
-C

**Figure S2. Identification of genuine inovirus pI proteins.** A. Characteristics of protein clusters (PCs) including pI-like ("Morphogenesis") proteins, i.e. proteins with a best hit to a pI-like model, and used as seed to identify inovirus sequences. Each PC is associated with a protein family (iPF), the number of proteins in the cluster, their initial affiliation, their origin (genome or metagenome), and host information for the ones identified in microbial genomes. B. Schematic representations of the different types of pI proteins identified: typical with an N-terminal Zot-like domain followed by a transmembrane domain (TMD), atypical with an N-terminal TMD followed by a Zot-like domain, and potential pI – Assembly fusions including an N-terminal Zot-like domain followed by a TMD and a secretion system-like domain.

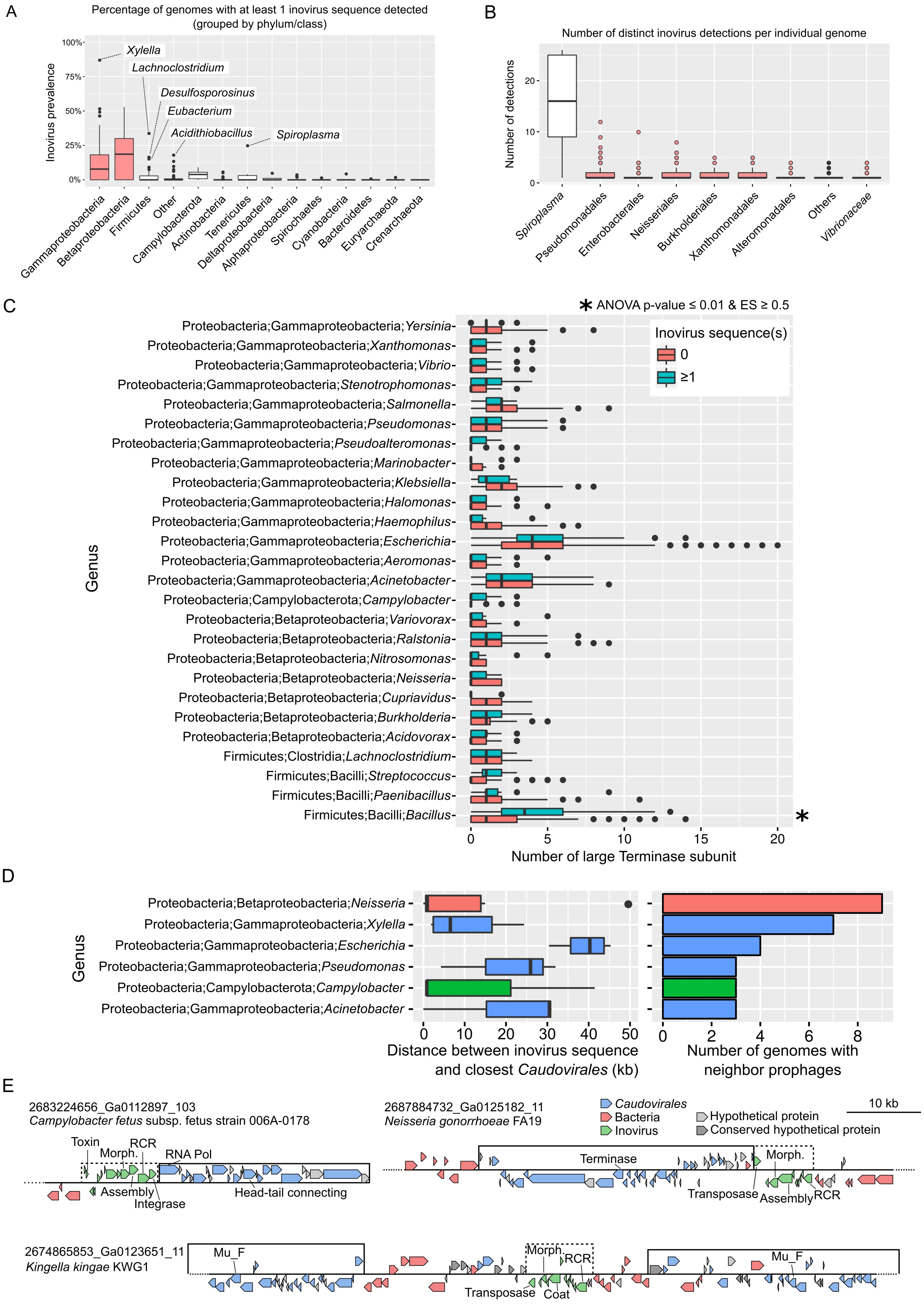

A
Percentage of genomes with at least 1 inovirus sequence detected (grouped by phylum/class)
Inovirus prevalence
Xylella
Lachnoclostridium
Desulfosporosinus
Eubacterium
Acidithiobacillus
Spiroplasma
Gammaproteobacteria
Betaproteobacteria
Firmicutes
Other
Campylobacterota
Actinobacteria
Tenericutes
Deltaproteobacteria
Alphaproteobacteria
Spirochaetes
Cyanobacteria
Bacteroidetes
Euryarchaeota
Crenarchaeota
B
Number of distinct inovirus detections per individual genome
Number of detections
Spiroplasma
Pseudomonadales
Enterobacterales
Neisseriales
Burkholderiales
Xanthomonadales
Alteromonadales
Others
Vibrionaceae
C
* ANOVA p-value ≤ 0.01 & ES ≥ 0.5
Inovirus sequence(s)
0
≥1
Genus
Proteobacteria;Gammaproteobacteria;Yersinia
Proteobacteria;Gammaproteobacteria;Xanthomonas
Proteobacteria;Gammaproteobacteria;Vibrio
Proteobacteria;Gammaproteobacteria;Stenotrophomonas
Proteobacteria;Gammaproteobacteria;Salmonella
Proteobacteria;Gammaproteobacteria;Pseudomonas
Proteobacteria;Gammaproteobacteria;Pseudoalteromonas
Proteobacteria;Gammaproteobacteria;Marinobacter
Proteobacteria;Gammaproteobacteria;Klebsiella
Proteobacteria;Gammaproteobacteria;Halomonas
Proteobacteria;Gammaproteobacteria;Haemophilus
Proteobacteria;Gammaproteobacteria;Escherichia
Proteobacteria;Gammaproteobacteria;Aeromonas
Proteobacteria;Gammaproteobacteria;Acinetobacter
Proteobacteria;Campylobacterota;Campylobacter
Proteobacteria;Betaproteobacteria;Variovorax
Proteobacteria;Betaproteobacteria;Ralstonia
Proteobacteria;Betaproteobacteria;Nitrosomonas
Proteobacteria;Betaproteobacteria;Neisseria
Proteobacteria;Betaproteobacteria;Cupriavidus
Proteobacteria;Betaproteobacteria;Burkholderia
Proteobacteria;Betaproteobacteria;Acidovorax
Firmicutes;Clostridia;Lachnoclostridium
Firmicutes;Bacilli;Streptococcus
Firmicutes;Bacilli;Paenibacillus
Firmicutes;Bacilli;Bacillus
Number of large Terminase subunit
D
Genus
Proteobacteria;Betaproteobacteria;Neisseria
Proteobacteria;Gammaproteobacteria;Xylella
Proteobacteria;Gammaproteobacteria;Escherichia
Proteobacteria;Gammaproteobacteria;Pseudomonas
Proteobacteria;Campylobacterota;Campylobacter
Proteobacteria;Gammaproteobacteria;Acinetobacter
Distance between inovirus sequence and closest Caudovirales (kb)
Number of genomes with neighbor prophages
E
2683224656_Ga0112897_103
Campylobacter fetus subsp. fetus strain 006A-0178
Toxin
Morph.
RCR
RNA Pol
Assembly
Integrase
Head-tail connecting
2687884732_Ga0125182_11
Neisseria gonorrhoeae FA19
Caudovirales
Bacteria
Inovirus
Hypothetical protein
Conserved hypothetical protein
10 kb
Terminase
Transposase
Assembly
RCR
2674865853_Ga0123651_11
Kingella kingae KWG1
Mu_F
Coat

**Figure S3. Inovirus prevalence and co-infection patterns.** A. Prevalence of inoviruses estimated through the proportion of genomes within a genus with 1 or more inovirus detection(s). Beta- and Gammaproteobacteria are highlighted in red. Groups with unusually high inovirus prevalence, > 75% within beta- or gamma-proteobacteria or > 15% otherwise, are labeled on the plot. B. Distribution of the number of distinct detection(s) by genome, grouped by host phylum or class. Host groups are colored as in panel A. For both boxplots, lower and upper hinges correspond to the first and third quartiles, whiskers extend no further than ±1.5*Inter-quartile range. C. Distribution of the number of large terminase subunits (TerL) as a proxy for the number of *Caudovirales* prophages identified by genome for each genus where ≥ 10 genomes had an inovirus detection and ≥ 10 genomes had no inovirus detection. Groups for which the distribution of prophage number was statistically different between the two categories (ANOVA p-value ≤ 0.01 & Cohen's effect size ≥ 0.5, degree of freedom=1) are highlighted with a star. D. Distribution of the distance between an inovirus prophage and the closest *Caudovirales* prophage for cases where the two sequences are less than 50kb apart. Distribution was plotted for genera where ≥ 3 cases of neighboring prophages were identified. Boxplot lower and upper hinges correspond to the first and third quartiles, whiskers extend no further than ±1.5*Inter-quartile range. Boxes are colored by host class. For panels A, B, C, and D, prevalence and co-infection frequencies were calculated after clustering near-clonal host genomes based on pairwise ANI (cutoffs: 95% nucleotide identity on 95% alignment fraction). E. Examples of (near-)contiguous inovirus and *Caudovirales* prophages. Three genome regions encoding both the inovirus and the *Caudovirales* prophages are displayed, with genes colored according to their affiliation. Prophages are highlighted with a solid black line (*Caudovirales*) or dashed black line (inovirus).

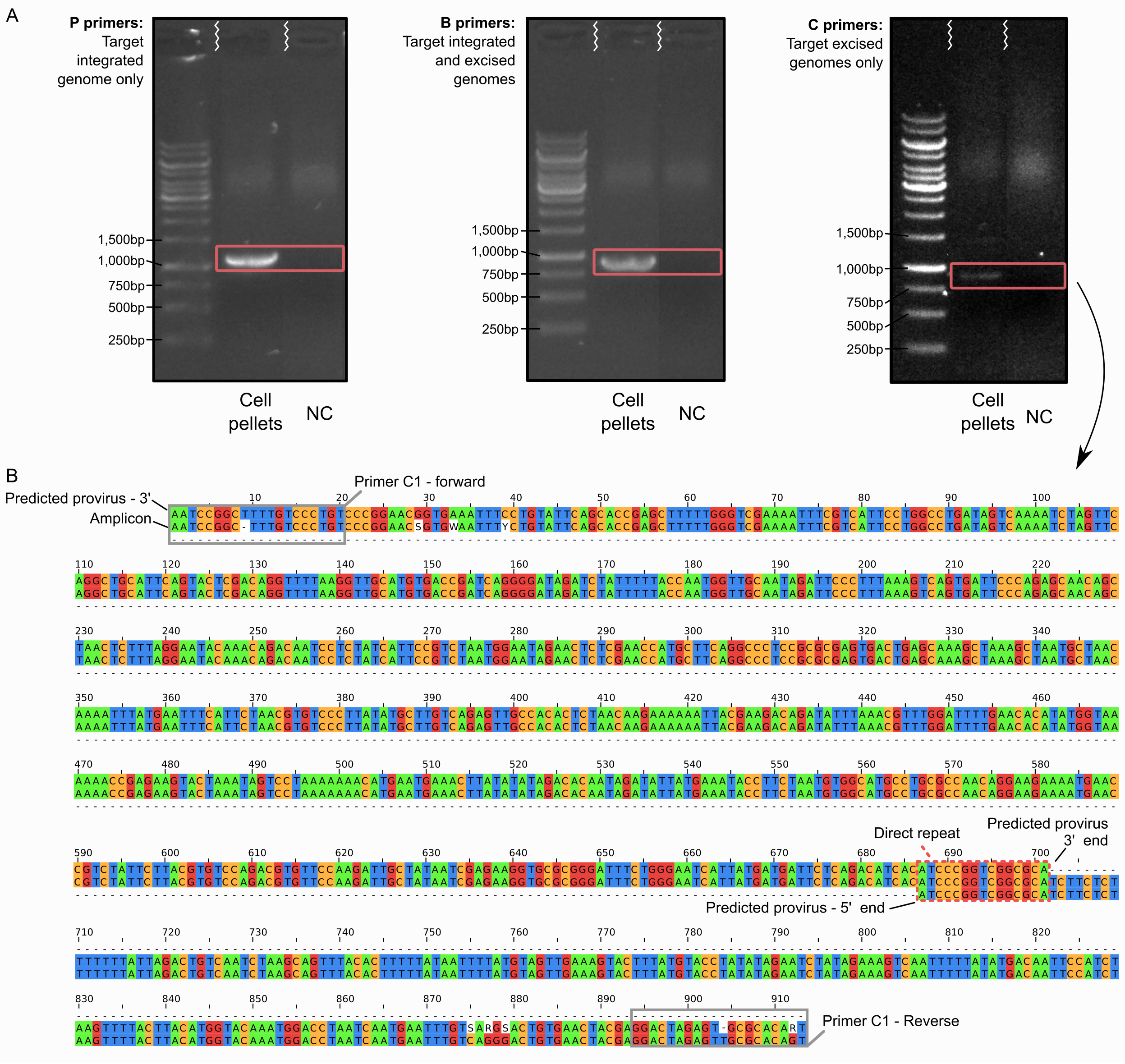


**Figure S4. Experimental validation of predicted provirus in *Methanolobus profundi* MobM.** A. Amplification result for the three primer pairs tested. P primers amplify across the predicted 5' insertion site (left), B primers amplify within the predicted provirus (center), and C primers amplify across the junction of the predicted excised circular genome (right). NC: no template control. B. Amplification products obtained with the C primer (i.e. spanning the junction of the predicted excised genome) aligned against the genome sequence of *Methanolobus profundi* MobM. Top track represents the 3' region of the provirus, bottom track the 5' region of the provirus, and the middle track is the sequenced amplicon. The direct repeat predicted as the end of the provirus is framed in red. Since the amplicon aligned across this direct repeat and from the 3' to the 5' end of the provirus, it is most likely derived from a circular excised version of the virus genome.

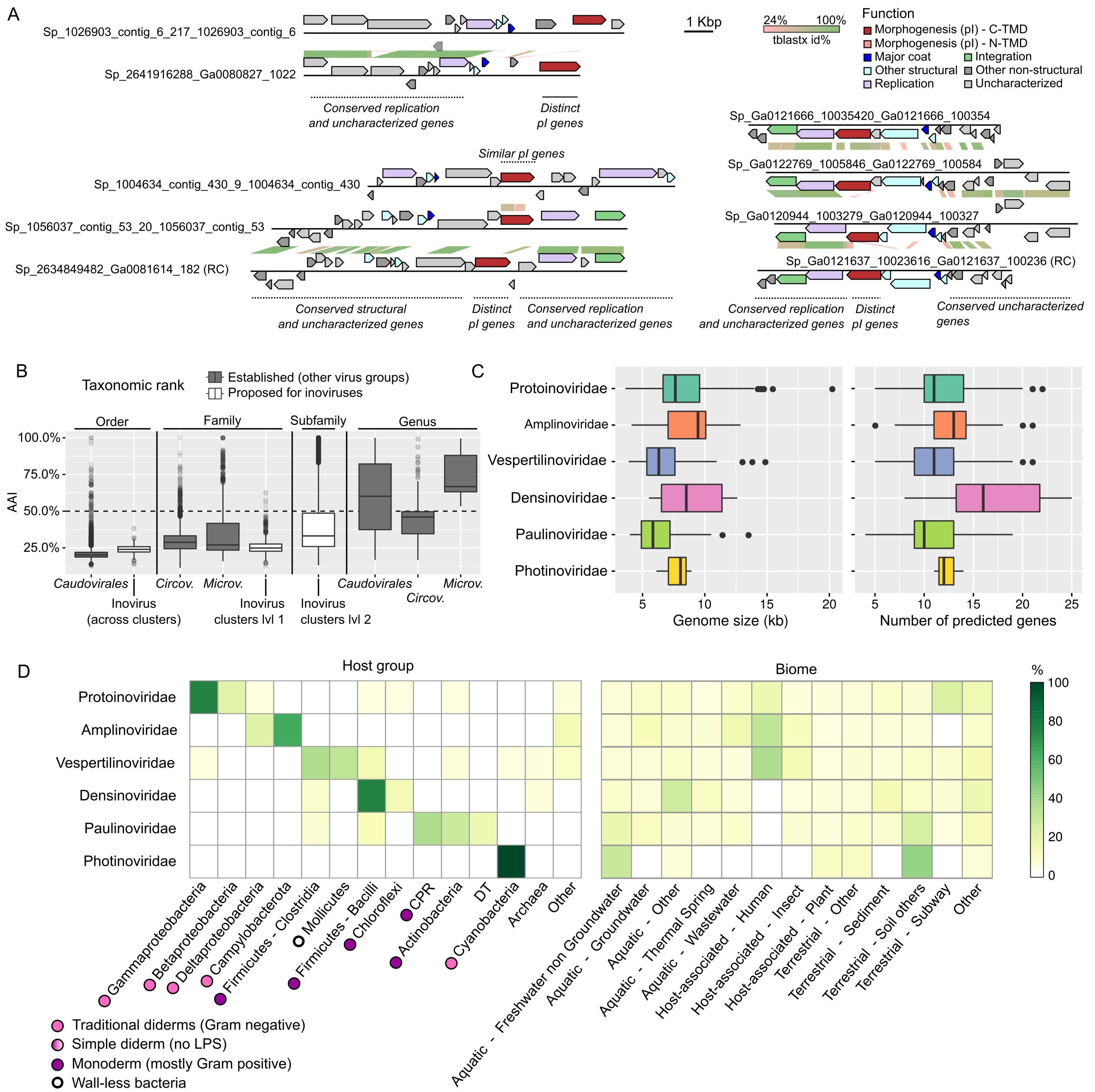


**Figure S5. Characteristics of the genome-based inovirus classification.** A. Examples of inovirus genomes with partial gene content sharing. Three comparisons of predicted inovirus genomes highlighting the fact that some of these viruses can display nearly-identical genes but show no similarity between morphogenesis (pI-like) proteins. Genes are colored according to their functional affiliation, based on the iPF clustering (Table S5). B. Distribution of pairwise marker gene Amino Acid Identity (AAI) for different viral groups and taxonomic ranks. Marker genes used included pI (Morphogenesis) for inoviruses, TerL (large terminase subunuit) for *Caudovirales*, Rep (replication initiation protein) for *Circoviridae*, and VP1 (major capsid protein) for *Microviridae*. Boxplots are colored according to the taxonomic ranks of the sequences compared. A dashed horizontal line indicates the threshold recently proposed to delineate *Inoviridae* genera (50% AAI). *Circov.*: *Circoviridae*, *Microv.*: *Microviridae*. Boxplot lower and upper hinges correspond to the first and third quartiles, whiskers extend no further than ±1.5* Inter-quartile range. C. Characteristic genome features of proposed families. Boxplots show the distribution of genome size (left) and number of predicted genes (righ) for each newly proposed family, colored as in Figure 5. Genome size and number of predicted genes were only calculated on inovirus genomes reliably predicted as complete, i.e. isolates, circular contigs, or proviruses with a confident insertion site either in a tRNA or next to an integrase gene. Boxplot lower and upper hinges correspond to the first and third quartiles, whiskers extend no further than ±1.5*Inter-quartile range. D. Host and biome range of proposed inovirus families. For each candidate family, the percentage of species associated with a specific host group (left) or ecosystem type (right) is indicated. Only host groups and biomes associated with > 10% of the species of at least 1 candidate family are indicated separately, the remaining are gathered in the "Other" category. Type of membrane for host cells are derived from ref.[23]. DT: Deinococcus-Thermus.

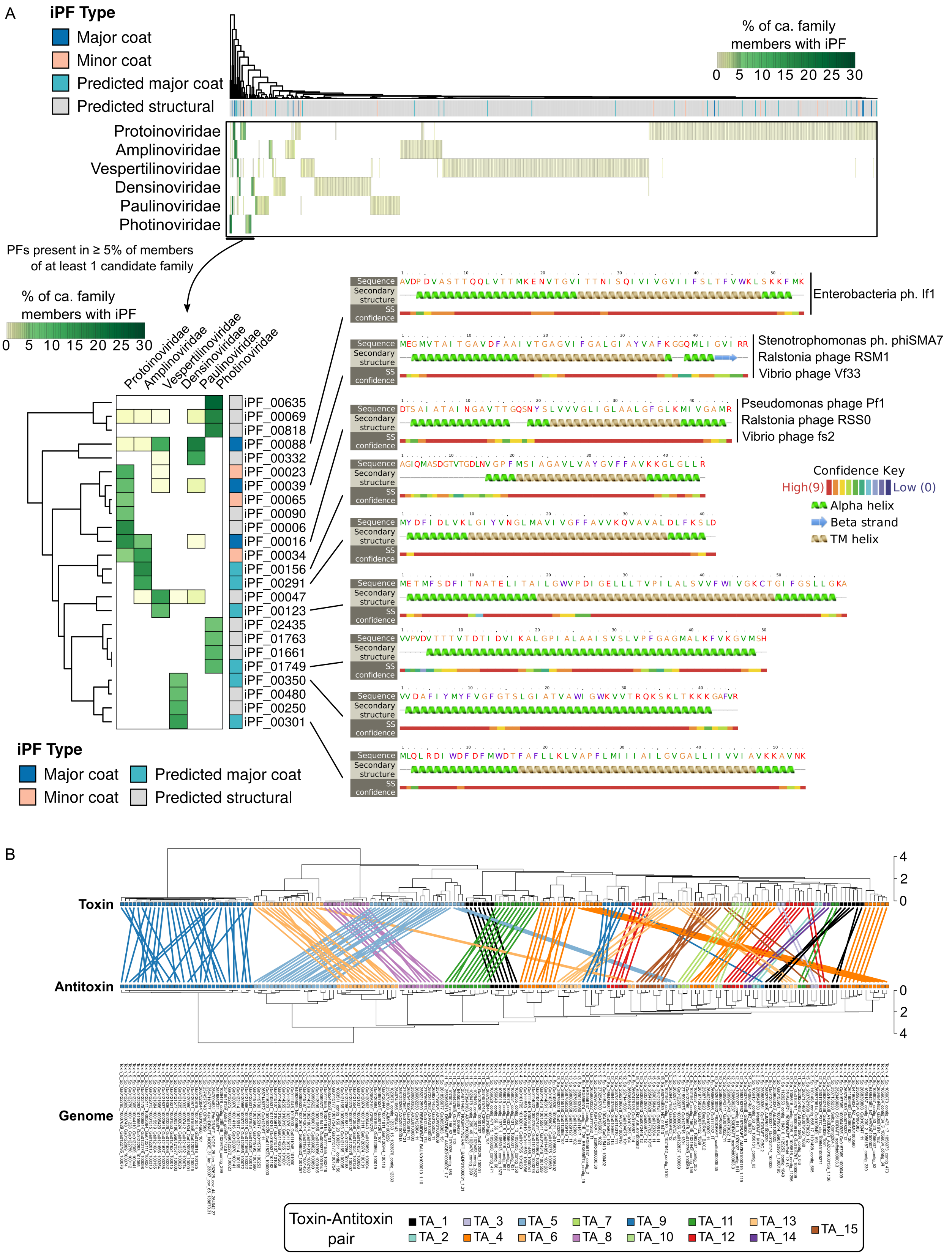


**Figure S6. Distribution of structural proteins and toxin/antitoxin proteins across inovirus species**. A. Diversity of structural proteins across inovirus sequences. The top heatmap displays
580 the relative abundance of each protein family (iPF) predicted as structural in each proposed

family. Color scale represents the percentage of members of the proposed family encoding each iPF. A zoomed heatmap displaying only iPFs found in ≥ 5% of members of ≥ 1 proposed family is displayed in the bottom left corner. Secondary structure predictions obtained from Phyre2[24] are displayed on the right side for the most abundant iPFs predicted as major coat proteins for each candidate family. B. Comparison of predicted toxin and antitoxin proteins similarities. Sequences predicted as toxins and antitoxins were compared using SDT, and the resulting AAI matrix was used to cluster sequences (UPGMA clustering). Predicted toxin-antitoxin (TA) pairs are highlighted with colors. The corresponding genome of the system is indicated at the bottom in the same order as the antitoxin gene.

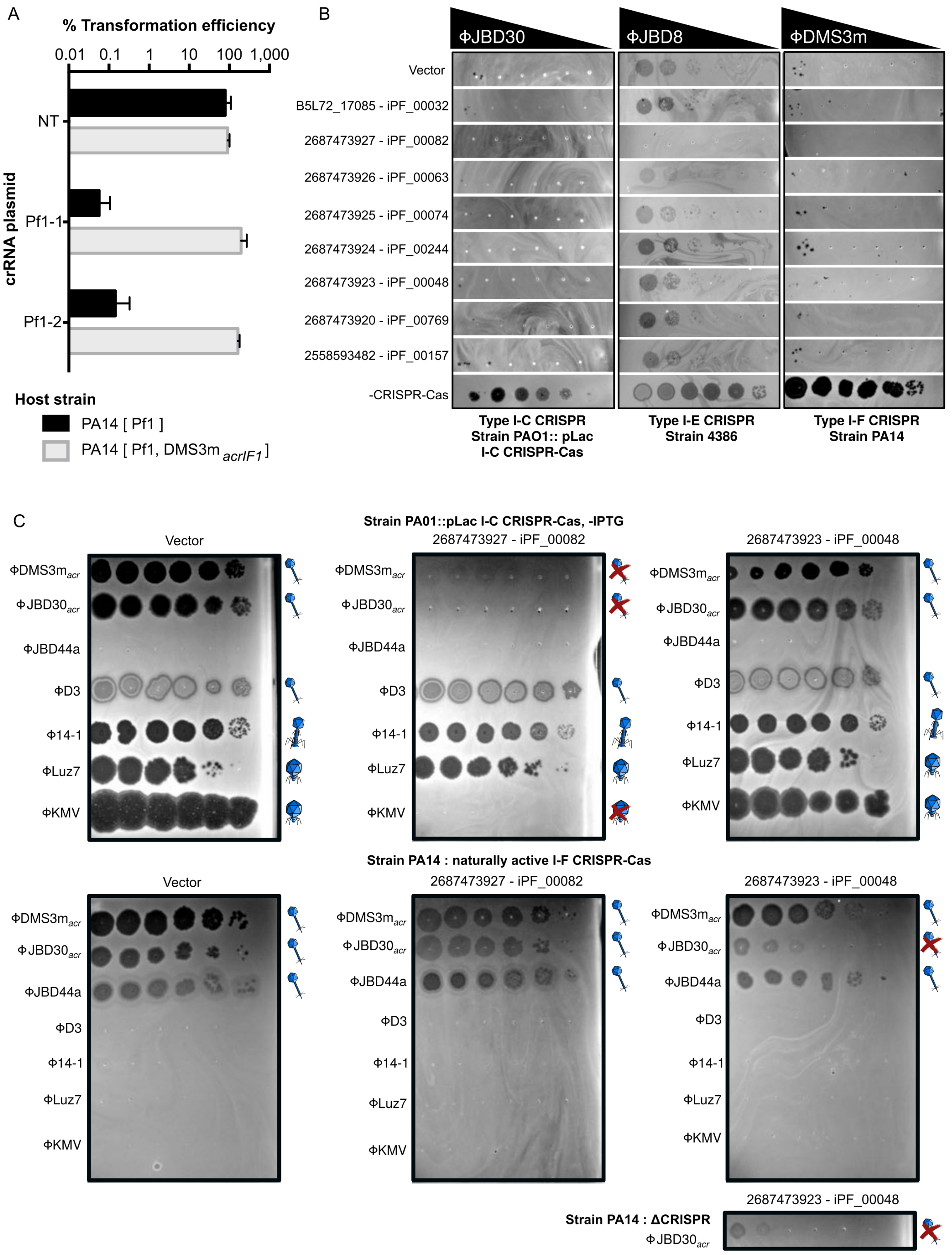


**Figure S7. Evaluation of self-targeting lethality, trans-acting anti-CRISPR activity from co-infecting prophages, and anti-CRISPR/superinfection activity of novel genes predicted on inovirus prophages in *Pseudomonas aeruginosa*.** A. Transformation assay to evaluate viability of cells including a self-targeted inovirus in the presence and absence of a co-infecting acr-encoding prophage. Percent transformation efficiency of crispr RNA (crRNA)-expressing plasmids were calculated relative to an empty vector, in *Pseudomonas aeruginosa* strains PA14 naturally lysogenized with inovirus Pf1 (PA14 [Pf1]) or dual lysogenized with Pf1 and acr-expressing siphovirus DMS3m$_{\mathit{acrIF1}}$ (PA14 [Pf1 , DMS3m$_{\mathrm{acrIF1}}$]). NT = non-targeting crRNA, Pf1-1 and Pf1-2 crRNAs target the coat protein gene in inovirus Pf1. B. Phage plaque assay to assess anti-CRISPR activity of candidate genes, using 3 host strains (left, middle, and right panel) each expressing a different type of CRISPR-Cas system, and the corresponding targeted phages (indicated on top of each panel). Host strains 4386 and PA14 encode a naturally active Type I-E and Type I-F CRISPR-Cas system (respectively), while strain PAO1 encodes Type I-C Cas genes integrated under the control of an IPTG inducible promoter, in presence of IPTG. Ten-fold serial dilutions of the targeted phages were titered on lawns of *Pseudomonas aeruginosa* expressing the empty vector (top row), a candidate gene (rows 2 to 11), or with CRISPR immunity suppressed (bottom row, condition -CRISPR-Cas). C. Phage plaque assays illustrating superinfection exclusion properties of genes 2687473927 (middle panel) and 2687473923 (right panel), relative to vector control (left panel). Serial dilutions (from left to right) of a set of phages (rows 1 to 7 in each picture) were spotted onto lawn cultures of strain PAO1 with the I-C Cas genes integrated under the control of an IPTG inducible promoter in the absence of IPTG (top), or of strain PA14 (bottom). Interpretation of infection outcome is indicated to the right of each lane, with successful infection represented by a phage symbol, and superinfection exclusion represented by a phage symbol barred by a red cross. To confirm that the inhibitory phenotype of 2687473923 on phage JBD30 and host PA14 is CRISPR-independent, the assay was repeated in a strain of PA14 lacking an active Type I-F system (PA14 ΔCRISPR, bottom right).

### Supplementary Tables

**Table S1. List and characteristics of reference inovirus genomes used in this study.** For each genome, genome features (size and type), ICTV classification, and known or predicted major coat proteins are indicated. Proteins that were not annotated as major coat but only predicted based on protein size and the presence of a single transmembrane domain (TMD) are highlighted in yellow. The tab "Structural protein detections" includes the detection of all putative structural proteins (i.e. major and minor coat proteins) in the same reference genomes. TMD: transmembrane domain.

**Table S2. List of genomes and metagenomes mined.** Genomes are associated with their IMG identifiers and taxonomic affiliation, with amendment to this affiliation specifically for the inovirus-encoding contigs added in the "Notes" column. Metagenomes are associated with their GOLD biome classifications, as well as the summarized ecosystem categories and subcategories used for Figure 2.

**Table S3. Classification of inovirus sequences into species, proposed families, and proposed subfamilies.** Putative tandem detections, i.e. neighboring inovirus prophages for which clear boundaries could not be identified, are shown in a separate tab ("Tandems") and were not included in the network from which the family/subfamily classification was derived. Each sequence is associated with its host genome affiliation or the sample ecosystem classification of the metagenome it was assembled from.

**Table S4. Additional indication of inovirus infection for 20 new phylum-level putative host groups.** Since inovirus sequences have only been detected in a (draft) genome for these groups, they could potentially originate from genome contamination, either physical sample contamination or *in silico* contamination for metagenome-assembled genomes. Two indicators were used to confirm the host linkage and alleviate this potential contamination: the presence of an integrated inovirus in a large host contig with confident affiliation, and the presence of match(es) between CRISPR spacer(s) and predicted inovirus sequence(s). These examples are listed here for each group highlighted in bold in Figure 3.

**Table S5. Functional annotation of protein families (iPFs).** Protein sequences were affiliated against the PFAM database and reference protein clusters derived from isolate inoviruses (affiliations starting with "PC_"). In the absence of significant hits to PFAM or the reference inovirus protein clusters, protein sequences predicted as putative structural proteins based on sequence characteristics were affiliated as "Predicted_structural", "Predicted_structural_SP", or "Putative_structural" depending on the prediction confidence (see Table S1, tab "Structural proteins detections"). iPFs were then organized in a two levels functional classification (columns 3 and 4). Identification of motifs for replication and integration iPFs as well as toxin-antitoxin pair iPFs are shown in separate tabs. Conserved domains were identified in iPFs affiliated as replication initiation and integration proteins, except for cases where too few sequences were available to reliably identify motifs (identified with "-"). Putative toxin-antitoxin are identified as pairs of co-occurring iPFs systematically located next to each other in inovirus genomes and for which at least one member of the pair was affiliated as either a putative toxin or antitoxin.

**Table S6. List of matches between inovirus sequences and IMG CRISPR spacer database.** Only cases with 0 or 1 mismatch between the spacer and putative viral sequences are included. Characteristics of host genomes with inovirus self-target, i.e. CRISPR spacer maching an integrated inovirus prophage in the same genome, are indicated in a separate tab. For each match, the prophage and spacer ID is indicated, along with the list of putative anti-CRISPR proteins, the detection of non-inovirus prophages in the same genomes (VirSorter predictions and identification of large terminase subunit), and the number of uncharacterized proteins with an HTH domain identified in the inovirus genome (using the representative genome from the inovirus species).

### References


1. Rosvall, M. & Bergstrom, C. T. Multilevel compression of random walks on networks reveals hierarchical organization in large integrated systems. *PLoS One* **6,** (2011).

2. Remmert, M., Biegert, A., Hauser, A. & Söding, J. HHblits: lightning-fast iterative protein sequence searching by HMM-HMM alignment. *Nat. Methods* **9,** 173–175 (2011).

3. Li, W. & Godzik, A. Cd-hit: A fast program for clustering and comparing large sets of protein or nucleotide sequences. *Bioinformatics* **22,** 1658–1659 (2006).

4. Edgar, R. C. MUSCLE: a multiple sequence alignment method with reduced time and space complexity. *BMC Bioinformatics* **5,** 113 (2004).

5. Eddy, S. R. Accelerated Profile HMM Searches. *PLoS Comput. Biol.* **7,** e1002195 (2011).

6. Fouts, D. E. Phage_Finder: automated identification and classification of prophage regions in complete bacterial genome sequences. *Nucleic Acids Res.* **34,** 5839–51 (2006).

7. Davis, B. M., Moyer, K. E., Fidelma Boyd, E. & Waldor, M. K. CTX prophages in classical biotype Vibrio cholerae: Functional phage genes but dysfunctional phage genomes. *J. Bacteriol.* **182,** 6992–6998 (2000).

8. Bille, E. *et al.* A virulence-associated filamentous bacteriophage of Neisseria meningitidis increases host-cell colonisation. *PLoS Pathog.* **13,** 1–23 (2017).

9. Ku, C., Lo, W. S., Chen, L. L. & Kuo, C. H. Complete genomes of two dipteran-associated spiroplasmas provided insights into the origin, dynamics, and impacts of viral invasion in Spiroplasma. *Genome Biol. Evol.* **5,** 1151–1164 (2013).

10. Roux, S., Hallam, S. J., Woyke, T. & Sullivan, M. B. Viral dark matter and virus-host interactions resolved from publicly available microbial genomes. *Elife* **4,** e08490 (2015).

11. Díaz-Muñoz, S. L., Sanjuán, R. & West, S. Sociovirology: Conflict, Cooperation, and Communication among Viruses. *Cell Host Microbe* **22,** 437–441 (2017).

12. Adriaenssens, E. M., Krupovic, M. & Knezevic, P. Taxonomy of prokaryotic viruses : 2016 update from the ICTV bacterial and archaeal viruses subcommittee. *Arch. Virol.* **162,** 1153–1157 (2017).

13. Iranzo, J., Krupovic, M. & Koonin, E. V. The double-stranded DNA virosphere as a modular hierarchical network of gene sharing. *MBio* **7,** e00978-16 (2016).

14. Wu, E. *et al.* Characterization of a cryptic plasmid from Bacillus sphaericus strain LP1-G. *Plasmid* **57,** 296–305 (2007).

15. Ilyina, T.V. ; Koonin, E. V., Ilyina, T. V & Koonin, E. V. Conserved sequence motifs in the initiator proteins for rolling circle DNA replication encoded by diverse replicaons

from eubacteria, eucaryotes and archaeabacteria. *Nucleic Acids Res.* **20,** 3279–3285 (1992).

16. Kimura, M., Wang, G., Nakayama, N. & Asakawa, S. in *Biocommunication in Soil Microorganisms.* (ed. Witzany, G.) 189–213 (Springer Berlin Heidelberg, 2011).

17. Wang, Y. *et al.* Identification, characterization, and application of the replicon region of the halophilic temperate sphaerolipovirus SNJ1. *J. Bacteriol.* **198,** 1952–1964 (2016).

18. Krupovic, M., Cvirkaite-Krupovic, V., Iranzo, J., Prangishvili, D. & Koonin, E. V. Viruses of archaea: Structural, functional, environmental and evolutionary genomics. *Virus Res.* **244,** 181–193 (2018).

19. Mochimaru, H. *et al.* Methanolobus profundi sp. nov., a methylotrophic methanogen isolated from deep subsurface sediments in a natural gas field. *Int. J. Syst. Evol. Microbiol.* **59,** 714–718 (2009).

20. Edwards, R. A., McNair, K., Faust, K., Raes, J. & Dutilh, B. E. Computational approaches to predict bacteriophage-host relationships. *FEMS Microbiol. Rev.* **40,** 258–272 (2016).

21. Kim, A. Y. & Blaschek, H. P. Construction and characterization of a phage-plasmid hybrid (phagemid), pCAK1, containing the replicative form of viruslike particle CAK1 isolated from Clostridium acetobutylicum NCIB 6444. *J. Bacteriol.* **175,** 3838–3843 (1993).

22. Bondy-Denomy, J. *et al.* Prophages mediate defense against phage infection through diverse mechanisms. *ISME J.* **22,** 1–13 (2016).

23. Gupta, R. S. Origin of diderm (Gram-negative) bacteria: Antibiotic selection pressure rather than endosymbiosis likely led to the evolution of bacterial cells with two membranes. *Antonie van Leeuwenhoek, Int. J. Gen. Mol. Microbiol.* **100,** 171–182 (2011).

24. Kelley, L. A., Mezulis, S., Yates, C., Wass, M. & Sternberg, M. The Phyre2 web portal for protein modelling, prediction, and analysis. *Nat. Protoc.* **10,** 845–858 (2015).
